# Interactions between cytoplasmic and nuclear genomes confer sex-specific effects on lifespan in *Drosophila melanogaster*

**DOI:** 10.1101/842716

**Authors:** Rebecca C. Vaught, Susanne Voigt, Ralph Dobler, David J. Clancy, Klaus Reinhardt, Damian K. Dowling

## Abstract

A large body of studies has demonstrated that genetic variation that resides outside of the cell nucleus can affect the organismal phenotype. The cytoplasm is home to the mitochondrial genome and, at least in arthropods, often hosts intracellular endosymbiotic bacteria such as *Wolbachia*. While numerous studies have implicated epistatic interactions between cytoplasmic and nuclear genetic variation as key to mediating patterns of phenotypic expression, two outstanding questions remain. Firstly, the relative contribution of mitochondrial genetic variation to other cytoplasmic sources of variation in shaping the phenotypic outcomes of cyto-nuclear interactions remains unknown. Secondly, it remains unclear whether the outcomes of cyto-nuclear interactions will manifest differently across the two sexes, as might be predicted given that cytoplasmic genomes are screened by natural selection only through females as a consequence of their maternal inheritance. Here, we address these questions, creating a fully-crossed set of replicated cyto-nuclear populations derived from three geographically distinct populations of *Drosophila melanogaster*, and measuring the lifespan of males and females from each population. We report cyto-nuclear interactions for lifespan, with the outcomes of these interactions differing across the sexes, and reconcile these findings with information on the full mitochondrial sequences and *Wolbachia* infection status of each of the populations.

## INTRODUCTION

Traditionally, most research investigating the genetic basis of variation in physiological and life-history trait expression in metazoans has focused on the role of genetic variation within the nuclear genome (Pesole et al., 2012). However, over the past two decades, it has become increasingly apparent that genetic elements that lie outside of the nuclear genome might commonly contribute to the expression of these traits. Such extra-nuclear genetic elements include the large community of microorganisms that colonise the host (such as in the digestive system), which collectively comprise the microbiome (Salvucci, 2016), several intracellular endosymbiotic bacteria, such as *Buchnera* (Moya et al., 2008), *Orientia* (Seong et al., 2001), and *Wolbachia* (Werren et al., 2008) that are commonly found in arthropods, as well as the genomes found within the mitochondria of virtually all contemporary eukaryotes, and the chloroplasts of plants. These extra-nuclear sources of genetic variation could in theory account for some of the documented “missing heritability” (Manolio et al., 2009) underpinning diseases and complex phenotypes, through either additive effects on the phenotype, or epistatic effects involving extra-nuclear and nuclear genetic variation.

In particular, research into the evolutionary consequences of extra-nuclear sources of genetic variation in arthropods has uncovered an unambiguous role for *Wolbachia* (Werren, 1997, 2008), and more recently mitochondrial genetic variation (Blier et al., 2001; Ballard & Whitlock, 2004; Ballard & Rand, 2005; Dowling et al., 2008; Burton et al., 2013; Dowling, 2014; Ballard & Pichaud, 2014; Hill et al., 2019), in contributing to trait expression. The *Wolbachia* endosymbiotic bacterium has been shown to exert influences on the biology of its hosts, ranging from pronounced effects on the phenotype such as cytoplasmic incompatibility (CI) in some arthropod species, to neutral or minor manipulations of cellular and reproductive processes; effects hypothesized to have evolved because they increase the relative fitness of *Wolbachia*-infected individuals within a population (Zug & Hammerstein, 2015; Correa & Ballard, 2016). *Wolbachia* has also been demonstrated to influence other host phenotypes such as lifespan and survival outcomes (Fry & Rand, 2002; Fry et al., 2004; Alexandrov et al., 2007; Maistrenko et al., 2016; Capobianco et al., 2018). Furthermore, other evidence has suggested that the magnitude of effects of *Wolbachia* infection on host phenotype can change across the nuclear genetic background of their hosts (Fry & Rand, 2002; Fry et al., 2004; Capobianco et al., 2018), implying that epistatic interactions between *Wolbachia* and their hosts may contribute to the host phenotypic expression (Correa & Ballard, 2016).

When it comes to the mitochondria, although the sequence variation within this genome was traditionally assumed to be evolutionarily neutral (Avise, 1986), many studies over the past quarter century have challenged this assumption (Ballard & Kreitman, 1994, 1995; Rand, 2001; Dowling et al., 2008; Hill et al., 2019). Mitochondrial genetic variation has been shown to affect the expression of numerous traits spanning physiology and life-history (reviewed in Ballard & Melvin, 2010; Horan et al., 2013; Wolff & Gemmell, 2013; Ballard & Pichaud, 2014; Vaught & Dowling, 2018). The exact mechanisms underlying how this variation has been able to accumulate have not been fully characterized. It is possible that this genetic variation is comprised predominantly of nonadaptive mutations, as a result of an inefficacy of selection in shaping the genome, and debate persists as to the relative level of purifying selection which acts on the mitochondrial genome (Popadin et al., 2012; Cooper et al., 2015; James et al., 2016; Shtolz & Mishmar, 2019). Alternatively, at least some of this genetic variation could comprise adaptive variants, and indeed several studies have recently provided evidence for this by demonstrating fitness benefits associated with particular mitochondrial variants within the environments in which they naturally exist (Blier et al., 2001; Rand, 2001; Mishmar et al., 2003; Ballard & Whitlock, 2004; Ruiz-Pesini et al., 2004; Meiklejohn et al., 2007; Ji et al., 2012; Dowling, 2014; Camus et al., 2017). Furthermore, given that the enzyme complexes that drive mitochondrial respiration are comprised of multiple polypeptide subunits encoded by both the mitochondrial genome and the nuclear genome (Taanman, 1999; McKenzie et al., 2007), and because many key elements of mitochondrial function are encoded by the nucleus (the genes regulating mtDNA replication, transcription, and translation machinery), a role for epistatic interactions between mitochondrial and nuclear genomes is likely to be important for maintaining cellular and ultimately organismal function (reviewed in Rand et al., 2004; Bar-Yaacov et al., 2012; Levin et al., 2014; Wolff et al., 2014; Stojković & Đordević, 2017; Hill et al., 2019; Shtolz & Mishmar, 2019).

Indeed, two bodies of empirical evidence indicate a role for mitochondrial-nuclear (mito-nuclear) interactions in shaping the evolutionary trajectories of populations. Firstly, several studies have reported evidence that nuclear-encoded proteins targeted to the mitochondria (N-mt proteins) have elevated rates of evolution relative to other nuclear-encoded proteins, presumably to accompany sequence changes within the mitochondrial genome, which exhibits a higher mutation rate than the nuclear genome (Barreto and Burton, 2013; Sloan et al., 2014; Havird et al., 2015; Sloan et al., 2017; Barreto et al., 2018; Yan et al., 2019). These co-associations between evolutionary rates of mitochondrial genes and nuclear-encoded mitochondrial genes suggest that these genes are coevolving. Indeed, several studies have documented signatures of linkage disequilibrium between polymorphisms across both nuclear and mitochondrial genomes in, for instance, humans, birds, flies, nematode worms and wild carrots (Jelić et al., 2015; Sloan et al., 2015; Haddad et al., 2018; Morales et al., 2018; Adineh & Ross, 2019; Cazares-Navarro & Ross, 2019; Ramsey et al., 2019; but see McKenzie et al., 2019). Notwithstanding, it remains to be determined whether the mode by which these genes are coevolving is predominantly compensatory (with selection acting on nuclear polymorphisms that restore negative effects imposed by mitochondrial mutations) or synergistic (selection for optimal combinations of mitochondrial-nuclear genotype) (Sloan et al., 2017; Hill et al., 2019). Secondly, several studies have harnessed experimental designs in which cytoplasmic genetic components from a set of lineages have been decoupled from their putatively coevolved (or “matched”) nuclear genetic backgrounds, and placed against novel (or “mismatched”) backgrounds of other lineages, to create panels of strains that possess either intra-lineage “matched”, or inter-lineage “mismatched” combinations of cytoplasmic and nuclear genotype. Such studies have been done by either mixing-and-matching cytoplasmic and nuclear genomes at an intra-specific (within or between populations of the same species) or interspecific (between species) scale. Such panels have been used to show that the genetic interactions between cytoplasm and nuclear genotype affect the expression of physiological and life-history traits (Wolff et al., 2014; Hill et al., 2019). This result has been substantiated by a recent meta-analysis across metazoan and plant kingdoms that reported that epistatic cyto-nuclear effects on trait expression tended to be larger than the additive cytoplasmic genetic effects (Dobler et al., 2014).

Furthermore, in studies that utilize cyto-nuclear strains, many of these have reported that the evolutionarily ‘mismatched’ combinations of mito-nuclear genotype (from different populations or even other species) exhibit lower fitness than the coevolved wild type combinations (Breeuwer & Werren, 1995; Kenyon & Moraes, 1997; Barrientos et al., 1998; Nagao et al., 1998; McKenzie et al., 2003; Roubertoux et al., 2003; Sackton et al., 2003; Zeyl et al., 2005; Ellison et al., 2008; Ellison & Burton, 2008; Lee et al., 2008; Chou et al., 2010; Osada & Akashi, 2012; Meiklejohn et al., 2013; Yee et al., 2013; Holmbeck et al., 2015; Zhu et al., 2015; Chang et al., 2016; Latorre-Pellicer et al., 2016; Ma et al., 2016; Jhuang et al., 2017; Tourmente et al., 2017; Healy & Burton, 2019; Pichaud et al., 2019), consistent with predictions that disruption of tightly coevolved mito-nuclear complexes would lead to fitness declines (Dowling, 2008; Lane, 2011; Hill et al., 2019). However, other studies have found that while the generation of novel combinations of mitochondrial and nuclear genotype typically led to altered expression of key traits, the fitness associated with the mismatched combinations was not consistently lower than the fitness of matched combinations (Rand et al., 2006; Dowling et al., 2007; Arnqvist et al., 2010; Montooth et al., 2010; Hoekstra et al., 2013; Zhu et al., 2014; Jelić et al., 2015; Immonen et al., 2016; Mossman et al., 2016a; Hoekstra et al., 2018; Dobler et al., 2018)

While our knowledge of the role of cyto-nuclear interactions in the regulation of organismal phenotype in metazoans has advanced considerably over the past decade (Hill et al., 2019), a full understanding of these interactions is currently limited by certain constraints. Firstly, several of the key studies conducted to date have focused on the consequences of combining cytoplasmic and nuclear genotypes sourced from populations that have exceptionally high levels of nucleotide divergence, or even from different species. This includes studies of the copepod *Tigriopus californicus*, a species that exhibits up to 20% divergence in mitochondrial sequences across populations (Barreto et al., 2018), and studies of the nematode *Caenorhabditis briggsae*, which has ∼2.6% divergence in mitochondrial protein coding genes across populations (Chang et al., 2016), as well as crosses involving mitochondrial and nuclear genotypes drawn from different sub-species or species in studies of mice and flies (Nagao et al., 1998; Roubertoux et al., 2003; Meiklejohn et al., 2013; Ma et al., 2016; Mossman et al., 2016a). While these studies provide proof-of-concept that cyto-nuclear interactions can modify phenotypic expression, inferences are drawn from crosses between distinct groups with high levels of mitochondrial genetic divergence, which may be unrepresentative of levels typically present across populations of most animal species. Moreover, studies that place cytoplasmic genomes from one species alongside highly divergent nuclear backgrounds drawn from distinct species, might inadvertently unmask cryptic mitochondrial genetic variation, which while adaptive or neutral when expressed alongside the pool of nuclear variation with which it has coevolved, could be non-adaptive and misaligned to optimal fitness when expressed alongside a highly novel nuclear background to which it has had no prior exposure (Chevin & Hoffmann, 2017; Dowling & Adrian, 2019).

Secondly, we have little understanding as to whether the outcomes of cyto-nuclear interactions manifest differently across males and females, since many studies of these interactions have previously focused on one sex only, or pooled the sexes in their analyses. The scope for the sex-specificity of such outcomes is particularly interesting in light of an evolutionary hypothesis called Mother’s Curse, which in its broadest form proposes that the maternal inheritance of cytoplasmic genetic elements, such as the mitochondrial genome and *Wolbachia* genome, will lead to the accumulation of variation that is neutral or beneficial to females, but deleterious to males (Cosmides & Tooby, 1981; Frank & Hurst, 1996; Gemmell et al., 2004; Havird et al., 2019). Extreme examples of this include the male-killing and the conversion of males into functional females caused by certain *Wolbachia* among arthropods (Werren et al., 2008). Although it was long thought that similar effects would not be applicable to the streamlined genome of the bilaterian metazoan mitochondria, a number of mitochondrial variants with male-harming effects, but with no obvious negative effects or even positive effects on female function, have been identified in flies, mice, hares, and humans (Nakada et al., 2006; Xu et al., 2008; Smith et al., 2010; Clancy et al., 2011; Innocenti et al., 2011; Camus et al., 2012; Patel et al., 2016; Aw et al., 2017; Milot et al., 2017; Camus & Dowling, 2018). The accumulation of sex-specific genetic variation in cytoplasmic genomes, particularly variation that is male-harming, may place selection on the host nuclear genome for modifier mutations that compensate for the cytoplasmically-induced effects (Turelli, 1994; Wade, 2014; Ågren et al., 2018; Connallon et al., 2018). This could then lead to sex differences in the trajectories of cyto-nuclear coevolution, potentially leading to greater levels of negative epistasis for fitness in males than females upon disruption of putatively coevolved combinations of cyto-nuclear genotype, if such disruption results in the unmasking of male-harming cytoplasmic genetic variants (Dowling & Adrian, 2019; Nagarajan-Radha et al., 2019 in-press). The results of recent studies that have partitioned cytoplasmic and nuclear genotypic contributions to the expression of adult life-history traits suggest that such a model of cyto-nuclear coevolution, based on nuclear modifiers rescuing the effects of male-harming cytoplasmic mutations, is plausible (Yee et al., 2013; Ma et al., 2016; Patel et al., 2016; Đorđević et al., 2015, 2017; Nagarajan-Radha et al., 2019 in-press). However, other studies investigating cyto-nuclear contributions to juvenile fitness traits and nuclear transcriptomic responses following inter-species cyto-nuclear mismatching have not found consistent male-biases (Mossman et al., 2016a, b; Mossman et al., 2017; Mossman et al., 2019).

And finally, in studies of invertebrates that utilize cyto-nuclear strains, cytoplasmic effects on the phenotype have typically been attributed to sequence variation in the mitochondrial genome, because in most cases the strains used were treated with antibiotics to eliminate concurrent infections with *Wolbachia* and other intracellular bacteria. However; the relative contributions of *Wolbachia* versus the mitochondrial genome to overall cytoplasmic effects on the phenotype currently remains unclear. Here, we further explore the capacity for cyto-nuclear interactions to confer sex-specific outcomes, by establishing a panel of fully replicated cyto-nuclear populations in *D. melanogaster*, in which we express cytoplasms from three geographically-distinct populations alongside representative pools of nuclear variation from each of the same three populations, in all nine possible combinations, each in triplicate. We then use these strains to investigate additive cytoplasmic, nuclear, and cyto-nuclear influences on lifespan of male and female individuals, to determine the capacity for cyto-nuclear interactions to manifest differently in each of the sexes, and explore the capacity for male-biases in negative epistasis upon mismatching of putatively coevolved cyto-nuclear gene complexes. We furthermore utilized a pooled-sequencing (pool-seq) approach to characterize the mitochondrial variation segregating within each of the cyto-nuclear populations, and determined the *Wolbachia* infection status of each, to enable us to explore the relative roles of *Wolbachia* infection, and mitochondrial variation, in contributing to observed cyto-nuclear interactions.

## MATERIALS AND METHODS

### Construction of cyto-nuclear populations

A set of fully-crossed cyto-nuclear populations of *D. melanogaster* was created (described also in Grunau *et al.,* (2018) and Dong *et al.,* (2019)) using three distinct laboratory populations. Each of these derived laboratory populations had originally been sourced from one of three separate continents. The first from Coffs Harbour, Australia (A) was originally collected in 2010 (Williams et al., 2012). The second population from Benin (B), Africa was originally collected in 1970, is routinely used for *Drosophila* genetics research, and is called the *Dahomey* population (Partridge and Andrews, 1985; Priest et al., 2007). The third population from Dundas, Canada (C) was originally collected in 2005 (MacLellan et al., 2009). Each of the three laboratory populations was initially founded by numerous mated-females collected from their respective wild populations, and was subsequently maintained at large population sizes thereafter under a 12:12 hour light: dark cycle at 25°C (Partridge & Andrews, 1985; Priest et al., 2007; MacLellan et al., 2009; Williams et al., 2012).

Prior to the creation of the cyto-nuclear populations used in our study, each of the three populations described above was maintained as a mass-bred population, kept across 10 vials, each propagated by 15 pairs of parents, at standardized egg densities of approximately 150 eggs per vial, developing on 6 mL of agar-based food substrate (containing potato, yeast, and dextrose), and maintained under a 12:12 hour light: dark cycle at 25°C. The offspring of each vial were admixed each generation, before adult individuals were redistributed into 10 vials of 15 pairs per vial. Upon receipt into our laboratory in 2010 and 2013, respectively, the Dahomey and Canadian populations were treated with the antibiotic tetracycline hydrochloride to remove *Wolbachia*. The Coffs Harbour population remained untreated.

Our goal was to create a set of cyto-nuclear populations that captured a large and representative sample of the segregating cytoplasmic and nuclear variation from each founding laboratory population, expressing each population-representative pool of cytoplasmic variation alongside pools of segregating nuclear variation from each of the founding populations, in all nine possible combinations (three cytoplasms [A,B,C] × three nuclear backgrounds [A,B,C]). That is, rather than sample one cytoplasmic variant and one nuclear variant per population, our design explicitly sought to preserve the pool of cytoplasmic and nuclear genetic variation segregating within each of the founding populations, within each cyto-nuclear population. This, therefore, enables us to screen for interactions between population-specific pools of cytoplasmic and nuclear genetic variation. Each of the nine cyto-nuclear combinations was created independently in triplicate, which increased our capacity to statistically partition effects attributable to cytoplasmic, nuclear and cyto-nuclear effects from confounding sources of variation across the population replicates.

To create the cyto-nuclear populations, 405 virgin females were collected from each of the three laboratory populations, and separated into groups of 45. Simultaneously, 405 males were collected from each of the same populations, and kept in groups of 45. Then, each group of 45 females was mixed with a group of males, in all possible intra- and inter-population combinations (3 × 3), with each cross combination replicated across three distinct replicates (thus, 27 population crosses in total). Each group of 45 pairs of individuals was kept across three vials of 15 pairs. Ten days later, the eclosing offspring of each of the three vials per combination were admixed together, and 45 virgin females collected from the admixed pool of offspring. These F_1_ females became the founders of a distinct cyto-nuclear population. Meanwhile, 405 males were collected from each of the three original source populations, and again separated into groups of 45. Each group of cyto-nuclear population females was then backcrossed to a group of males that came from the same population as the sires of these females, again across three vials of 15 pairs. In the next generations, the process was again repeated. This backcrossing procedure ran for 17 generations, resulting in nine cyto-nuclear combinations, each represented in three independent replicates (Supporting Information Fig. 1). In theory, 17 generations of backcrossing will have resulted in the replacement of more than 99.99% of the original nuclear alleles associated with each cytoplasmic background, thus effectively placing each cytoplasm alongside the nuclear variation from the target population. The backcrossing procedure was stopped two generations prior to the experiments commencing, to enable us to expand each of the cyto-nuclear populations to numbers required to set up the experiment described below.

### Lifespan assay

We assayed the adult lifespan of males and females from each cyto-nuclear population. The experiment was conducted across six sequential experimental sampling ‘blocks’, which were each separated in time by one generation. Males and females from each replicate cyto-nuclear population were assayed for lifespan in separate cohorts, in groups of 16 focal individuals per vial. Within each block, we assayed two cohorts of each sex for each replicate cyto-nuclear population (9 populations × 3 replicates × 2 sexes × 2 cohorts). Each cohort of focal individuals was placed with a group of 16 “tester” individuals of the opposite sex within a given experimental vial, such that each experimental vial housed 32 individuals in total. These mixed-sex conditions of even sex-ratio were employed with the aim of mimicking normal culture conditions for laboratory-adapted populations of *D. melanogaster* (Kimber & Chippendale, 2013). Tester individuals were collected from a laboratory-adapted outbred population, LHM, which was originally sourced from California, U.S.A. (Rice et al., 2005). These tester individuals carried an autosomal recessive mutation, which had been introgressed into the population, which encodes brown-eyes (LH_M-bw_) (Friberg & Dowling, 2008), distinguishing them from the wild type focal individuals.

The experiment described below was implemented according to an identical protocol across each experimental block. The only difference across blocks lay in the number of generations that had elapsed since the last generation of backcrossing (three generations in Block 1, incrementing to eight generations by Block 6). Experimental individuals and tester individuals were collected one day (at least 24 hours) after eclosion, meaning that most, if not all, individuals would have already mated during this time and were no longer virgins. All experimental individuals were collected under light CO_2_ anaesthesia from vials trimmed to approximately 150 eggs, laid by females that were five days old at the time of ovipositioning to mitigate maternal age effects (grandparental generations were also propagated by females that were five days of age at the time of ovipositioning and vials were also standardized for egg densities). Each vial used in the experiment contained a standard potato-dextrose food medium, and held 16 experimental individuals belonging to one sex and 16 tester individuals of opposite sex that were of identical age.

Once each of these experimental vials was established, each cohort was transferred to new vials with fresh food every second day, at which time the number of individuals that had died during the preceding 48 hours was scored. Expired individuals stuck to old food medium were scored for sex and eye-color, while individuals that were near-death (incapacitated but nonetheless moving) were carefully transferred to the corresponding new vial. To ensure a constant number of adult individuals (hence density) per vial, and an even sex ratio, all expired individuals (both focal individuals and tester individuals) were replaced with brown-eyed tester individuals of the same sex and similar age (within two weeks) from the LH_M-bw_ population every six days, for the first six weeks of each assay (Kimber & Chippendale, 2013).

### Mitochondrial genotyping of cyto-nuclear population

The mitochondrial genomes of the cyto-nuclear populations were sequenced using a pool-seq approach, sampling 150 individuals (sexes combined) from each of the 27 population replicates. DNA was extracted by first performing a mitochondrial enrichment protocol, where the individuals were first homogenized in isolation buffer (10 mM Tris-HCL, 5 mM EDTA, 250 mM sucrose, 0.2% BSA) using a 15 mL Dounce homogeniser on ice by eight strokes with the narrow gap pistil. The resulting supernatant was then filtrated and the homogenate was then centrifuged at 300 g for 3 minutes at 4° C. To precipitate the mitochondria, the supernatant was again filtrated and a subsequent centrifugation was performed for 10 minutes at 9000 g at 4° C. The resulting pellet was resuspended in 50 μL of isolation buffer. Once mitochondria were precipitated, mtDNA was extracted using the Analytic Jena innuPREP DNA Mini Kit (protocol 4), and the extracted mtDNA stored in elution buffer (Tris, pH=8). The extraction yielded between 15.5 μg/mL and 40.1 μg/mL of mtDNA for each line. The mtDNA was sequenced at a concentration of 400 ng/50 μL on an Illumina HiSeq platform using the ‘rapid run’ mode. The read length was 75 bp paired ends (PE) and resulted in an average 3.05 million reads for each sample (∼150x mean sequencing depth for each line). Prior to mapping, reads were trimmed to remove low quality base calls with a minimum quality score of 20 and minimum read length of 50. Reads were then mapped to the mitochondrial reference genome (NC_024511.2) using *BWA* v0.7.15-2 (Li, 2013). Unmapped reads were subjected to a second round of mapping using *Stampy* v1.6.30 (Lunter & Goodson, 2011). Subsequent quality filtering included removing reads with a mapping quality of less than 20 and duplicates. In order to detect significantly differentiated single nucleotide polymorphisms (SNPs) between the mitochondrial genomes from the cyto-nuclear populations (Australia, Benin, and Canada), the fixation index, FST (Hudson et al., 1992), and Fisher’s exact test for allele frequency differences were calculated per SNP in *PoPoolation2* v1.201 (Kofler et al., 2011) with a minimum depth of 30 and a minor allele frequency of 0.01. Those SNPs that were significant under Fisher’s exact test at a false discovery rate (FDR) (Benjamini & Hochberg, 1995) of 0.1%, were considered to be significantly differentiated.

### Wolbachia screening

A screen for *Wolbachia* infection was performed across each of the cyto-nuclear population replicates using the approach described in Richardson *et al.,* (2012) and Grunau *et al.,* (2018), involving diagnostic PCR, on DNA extracted from pools of 15 adult flies, using *Wolbachia*-specific *wsp* primer-sets (Richardson et al., 2012). This screen revealed that all populations harbouring Australian cytoplasms were infected with *Wolbachia*, while those possessing cytoplasms originating from Benin or Canada were uninfected.

### Statistical analysis

#### Partitioning of cytoplasmic and nuclear sources of variance

The data analyses were performed in the R statistical environment (v3.3.3, R core team). We first analysed whether lifespan was affected by the origin of each cytoplasm (3 levels; Australia, Benin and Canada), nuclear genetic background (3 levels, Australia, Benin and Canada), sex (males and females), or their interaction. We used a linear mixed effects model in the *lme4* package (Bates et al., 2014). The response variable was individual lifespan (measured in days), with cytoplasmic origin, nuclear background, and sex included as fixed factors, as well as interactions between the three. Random effects, which described the hierarchical structure of the data, were also included in the model. These consisted of experimental sampling block (n = 6), the vial identity in which the assays took place (n = 645), and the cyto-nuclear population replicate (n = 27). A full model was built with all the fixed effects, and including higher-order interactions involving fixed and random effects up to 2^nd^ order interactions. This full model was then reduced, by eliminating random interactions from the model that accounted for near-zero levels of variance in the model whose removal returned a non-significant *p* value indicated by Log-Likelihood Ratio testing using the *anova* function in R. Parameter values of fixed effects and their significance were estimated in the final model, using Type III Wald Chi-square tests in the *car* package (Fox, 2012) and random effects were estimated using the *summary* function..

We also conducted survival analyses using mixed model Cox proportional hazard models, modelling effects of the cytoplasm (3 levels: Australia, Benin and Canada), nuclear genetic background (3 levels, Australia, Benin and Canada), sex (males and females), and their interaction. These analyses are presented in the Supporting Information (Supporting Information Table 1) and are consistent with our interpretations based on the lifespan analyses described above.

#### Effects of cyto-nuclear mismatching

We ran a second model to explicitly test whether disruption of putatively coevolved cytoplasmic and nuclear genomes led to reduced lifespan, and if so, whether any such effects were sex-biased in magnitude or direction. To this end, we included a new factor, denoted coevolutionary status of the individual (‘matched’ or ‘mismatched’ cyto-nuclear genotype), and modelled lifespan, with coevolutionary status, sex and their interactions as fixed effects. The cytoplasm of origin, nuclear background, their interaction (3 x 3), as well as the replicate cyto-nuclear population (27 levels), experimental sampling block, and vial identity were included as random effects. Once the full model was built the same model reduction process, as described above, using Log-Likelihood ratio tests in order to reduce the fixed and random effects statement, was carried out to derive at a final reduced model. This reduced model returned a model convergence warning from R. In order to facilitate model convergence, a new model was constructed in which the parameter estimates for the fixed and random effects given by the original non-converging model were assigned as the starting values for the numerical algorithm for the new model with a set maximum of 20,000 iterations. Here, we present the results of the new converging model but note that the results of these two models (converging and non-converging) were virtually identical.

#### Homing in on mitochondrial genetic effects

The *Wolbachia* screen revealed that populations harbouring Australian cytoplasms were infected with *Wolbachia*, whereas other populations were uninfected. Furthermore, the genotyping analysis of the cyto-nuclear populations, which is reported in full in the Results, revealed that a small number of population replicates of particular cytoplasms, specifically AA3, CA1 and CA3 (notation is: Australian cytoplasm in Australian nuclear background, replicate 3; Canadian cytoplasm in Australian nuclear background, replicate 1; and Canadian cytoplasm in Australian nuclear background, replicate 3), were carrying dissimilar mitotypes (nearly fixed for different mtDNA SNPs) to the other population replicates of the same cytoplasm. In light of this, we conducted a reanalysis of the model outlined in the analysis above, excluding these outlier population replicates from the analysis. This analysis, where the outliers have been excluded, is presented in the Supporting Information (Supporting Information Table 2, Supporting Information Figure 2), and the results are qualitatively consistent with the results of the model containing all cytonuclear populations, which we present in the Results section.

#### Partitioning mitochondrial from *Wolbachia* effects

Finally, we explored the relative contributions of mitochondrial variation and heterogeneity in *Wolbachia* infection status in shaping the lifespan phenotype. All cyto-nuclear populations carrying the Australian cytoplasm were infected with *Wolbachia*, and thus it is challenging to determine whether effects attributable to this cytoplasm are caused by *Wolbachia* infection, mitochondrial genotype, or a combination of both. However, as mentioned above and outlined in full in the Results, one of the population replicates of the Australian cytoplasm in the Australian nuclear background (AA3), is fixed for an alternative mitotype compared to the other replicates of this same cyto-nuclear genotype (AA1 and AA2), and this provided the opportunity to explore whether lifespan associated with this *Wolbachia*-infected cytoplasm differed from that of the other *Wolbachia*-infected cytoplasms when expressed in the same Australian nuclear background (AA3 *versus* AA1 and AA2). Such a difference would point to a genuine mtDNA genotypic effect. Moreover, we compared lifespans of these three population replicates (AA1, AA2, AA3) to those of the Canadian and Benin mitotypes (which are *Wolbachia* free) when expressed in the same Australian nuclear background (CA and BA replicates). A general difference between all AA replicates and the CA and BA replicates would point to a key role for *Wolbachia* in driving the phenotypic effects observed in this study.

Specifically, within the Australian nuclear background, there are nine population replicates: three carry the Beninese cytoplasm (BA1, BA2, BA3), and as outlined in full in the Results section, two carry an Australian mitotype which is dominated by a haplotype previously denoted B1 (AA1, AA2), while one carries an Australian mitotype which is dominated by a haplotype previously denoted A1 (AA3) (Camus et al., 2017), one carries the Canadian mitotype (CA2), and CA1 and CA3 were included as their own separate grouping since they carry a mitotype seemingly unique from CA2 (See Results). Thus, the fixed effects in this model included mitotype (5 levels: A1, B1, CA1&CA3, CA2, and Benin), and sex (2 levels: males, females), and the random effects describing the hierarchical structure of the data incorporated cytoplasmic population (9 populations), sampling block (6 levels), and vial identity (218 levels). As described above, a full model was built with all the fixed effects, including higher-order interactions between fixed and random effects. This full model was then reduced, by eliminating fixed and random interactions from the model that accounted for near-zero levels of variance in the model, guided by Log-Likelihood ratio tests. A caveat exists for the analysis of this dataset. There is no independent population replicate of the AA3 mitotype, since this was only found in one of the three Australian cytoplasm × Australian nuclear replicates (AA3). Furthermore, there is no independent population replicate of the Canadian CA2 either, since the other two population replicates for this nuclear background (CA1 and CA3) were grouped separately because the mtDNA of these populations were more closely aligned to the mitotype present with the populations harbouring Beninese cytoplasms.

## RESULTS

### Mitochondrial genotyping of cyto-nuclear populations

We observed 202 SNPs in the mtDNA sequences of the populations in total, of which 34 were significantly differentiated (Fisher’s exact test at an FDR of 0.1%) between the cyto-nuclear populations (Table 1). These 34 SNPs included synonymous and non-synonymous changes, as well as nucleotide changes in non-coding RNAs and the origin of replication (Table 2). The majority of SNPs were significantly differentiated between the mtDNA sampled from the Australian laboratory population, and those of the other two laboratory populations (Benin and Canada). Genotype networks were also created based on the 34 highly differentiated SNPs and consensus sequences of the 27 replicate cyto-nuclear populations (Fig. 1). The consensus sequence of each population was defined by the most frequent nucleotide at each SNP site. Because a pool-seq approach was utilized, it is not possible to directly determine specific haplotypes. However, population-specific allele frequencies of the 34 significantly differentiated SNPs were generally high (Supporting Information Fig. 3-5), making it highly probable that high-frequency alleles, within each line, were found on the same haplotype.

**Figure 1.**
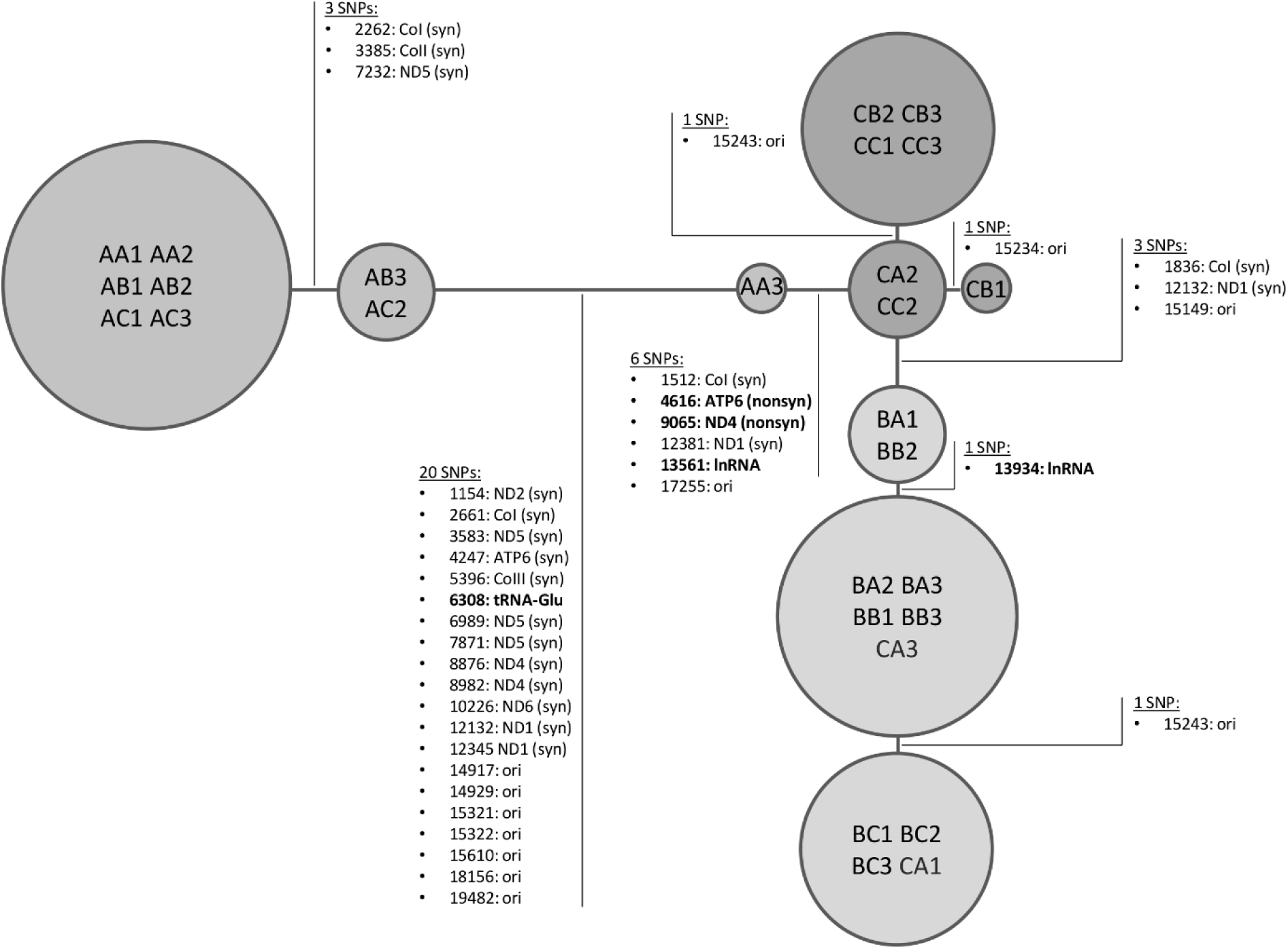
Network of mitochondrial genotypes of the 3 x 3 cyto-nuclear lines derived from the 34 significantly differentiated SNPs. The network is based on the consensus sequences of the lines. The consensus sequence of each population is defined by the most frequent nucleotide at each SNP site. Note that SNPs at positions 15,234 and 15,243 are of intermediate allele frequencies and differentiation at these two sites is relatively low (see also Table 1).

**Table 1.**
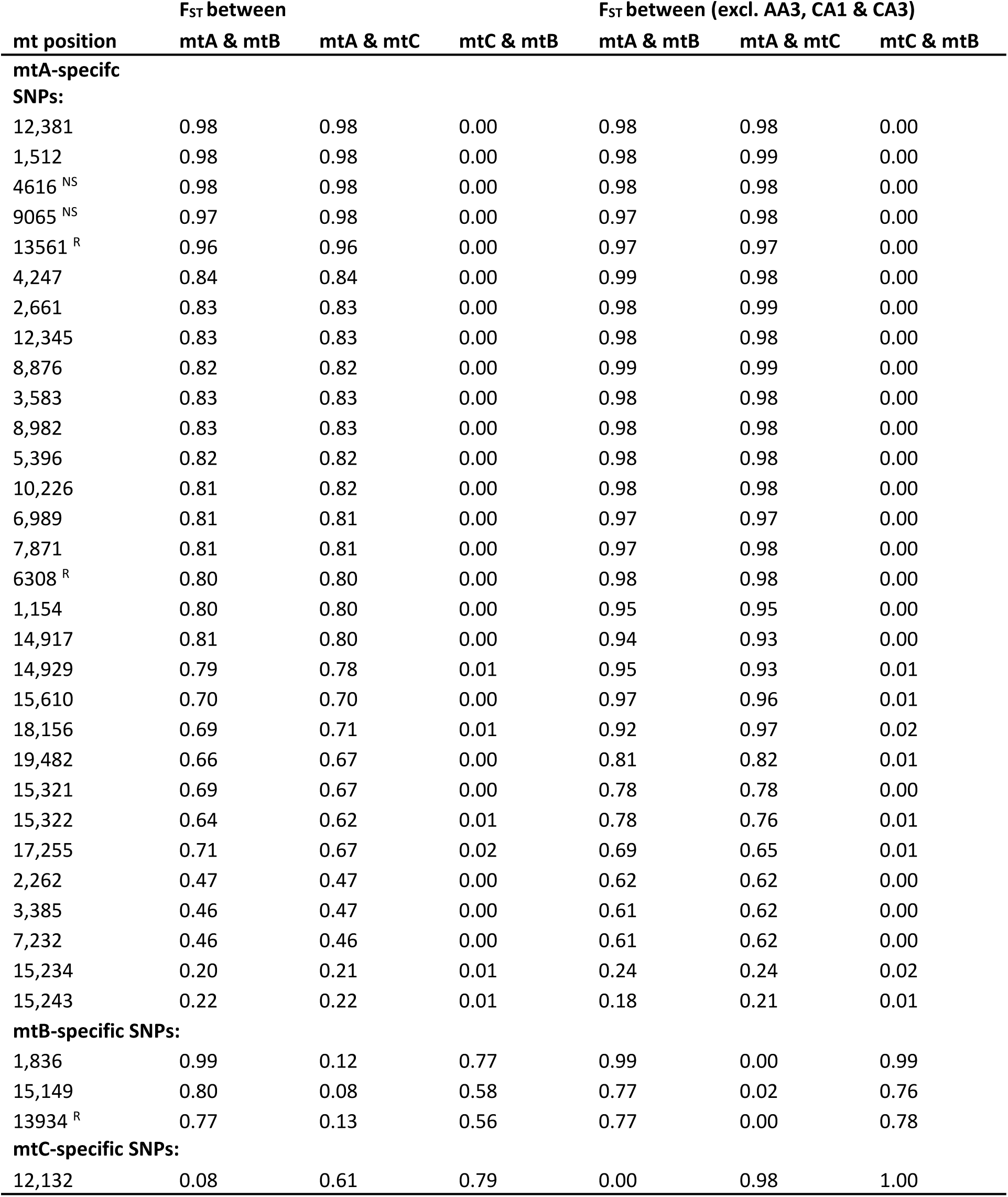
Mean pairwise FST of significantly differentiated mitochondrial single nucleotide polymorphisms (SNPs) (Fisher’s exact test; FDR=0.1%). SNPs are ranked according to FST. Pairwise FST between lines (replicates were pooled) was calculated based on Hudson *et al.,* (1992) using *PoPoolation2* (Kofler et al., 2011). Means of all pairwise FST between the mitochondrial genotypes derived from each population (Australia:A, Benin:B, Canada:C). Population replicates carrying outlier mitotypes AA3, CA1 and CA3 were excluded. Superscript lettering corresponds to nucleotide changes that result in non-synonymous (^NS^) amino acid changes and changes in non-coding RNAs (^R^). Mitochondrial (mt) position is according to mitochondrial reference sequence NC_024511.

| mt position | $F_{ST}$ between | | | $F_{ST}$ between (excl. AA3, CA1 & CA3) | | |
| --- | --- | --- | --- | --- | --- | --- |
|  | mtA & mtB | mtA & mtC | mtC & mtB | mtA & mtB | mtA & mtC | mtC & mtB |
| **mtA-specific SNPs:** |  |  |  |  |  |  |
| 12,381 | 0.98 | 0.98 | 0.00 | 0.98 | 0.98 | 0.00 |
| 1,512 | 0.98 | 0.98 | 0.00 | 0.98 | 0.99 | 0.00 |
| 4616<sup>NS</sup> | 0.98 | 0.98 | 0.00 | 0.98 | 0.98 | 0.00 |
| 9065<sup>NS</sup> | 0.97 | 0.98 | 0.00 | 0.97 | 0.98 | 0.00 |
| 13561<sup>R</sup> | 0.96 | 0.96 | 0.00 | 0.97 | 0.97 | 0.00 |
| 4,247 | 0.84 | 0.84 | 0.00 | 0.99 | 0.98 | 0.00 |
| 2,661 | 0.83 | 0.83 | 0.00 | 0.98 | 0.99 | 0.00 |
| 12,345 | 0.83 | 0.83 | 0.00 | 0.98 | 0.98 | 0.00 |
| 8,876 | 0.82 | 0.82 | 0.00 | 0.99 | 0.99 | 0.00 |
| 3,583 | 0.83 | 0.83 | 0.00 | 0.98 | 0.98 | 0.00 |
| 8,982 | 0.83 | 0.83 | 0.00 | 0.98 | 0.98 | 0.00 |
| 5,396 | 0.82 | 0.82 | 0.00 | 0.98 | 0.98 | 0.00 |
| 10,226 | 0.81 | 0.82 | 0.00 | 0.98 | 0.98 | 0.00 |
| 6,989 | 0.81 | 0.81 | 0.00 | 0.97 | 0.97 | 0.00 |
| 7,871 | 0.81 | 0.81 | 0.00 | 0.97 | 0.98 | 0.00 |
| 6308<sup>R</sup> | 0.80 | 0.80 | 0.00 | 0.98 | 0.98 | 0.00 |
| 1,154 | 0.80 | 0.80 | 0.00 | 0.95 | 0.95 | 0.00 |
| 14,917 | 0.81 | 0.80 | 0.00 | 0.94 | 0.93 | 0.00 |
| 14,929 | 0.79 | 0.78 | 0.01 | 0.95 | 0.93 | 0.01 |
| 15,610 | 0.70 | 0.70 | 0.00 | 0.97 | 0.96 | 0.01 |
| 18,156 | 0.69 | 0.71 | 0.01 | 0.92 | 0.97 | 0.02 |
| 19,482 | 0.66 | 0.67 | 0.00 | 0.81 | 0.82 | 0.01 |
| 15,321 | 0.69 | 0.67 | 0.00 | 0.78 | 0.78 | 0.00 |
| 15,322 | 0.64 | 0.62 | 0.01 | 0.78 | 0.76 | 0.01 |
| 17,255 | 0.71 | 0.67 | 0.02 | 0.69 | 0.65 | 0.01 |
| 2,262 | 0.47 | 0.47 | 0.00 | 0.62 | 0.62 | 0.00 |
| 3,385 | 0.46 | 0.47 | 0.00 | 0.61 | 0.62 | 0.00 |
| 7,232 | 0.46 | 0.46 | 0.00 | 0.61 | 0.62 | 0.00 |
| 15,234 | 0.20 | 0.21 | 0.01 | 0.24 | 0.24 | 0.02 |
| 15,243 | 0.22 | 0.22 | 0.01 | 0.18 | 0.21 | 0.01 |
| **mtB-specific SNPs:** |  |  |  |  |  |  |
| 1,836 | 0.99 | 0.12 | 0.77 | 0.99 | 0.00 | 0.99 |
| 15,149 | 0.80 | 0.08 | 0.58 | 0.77 | 0.02 | 0.76 |
| 13934<sup>R</sup> | 0.77 | 0.13 | 0.56 | 0.77 | 0.00 | 0.78 |
| **mtC-specific SNPs:** |  |  |  |  |  |  |
| 12,132 | 0.08 | 0.61 | 0.79 | 0.00 | 0.98 | 1.00 |

**Table 2.**
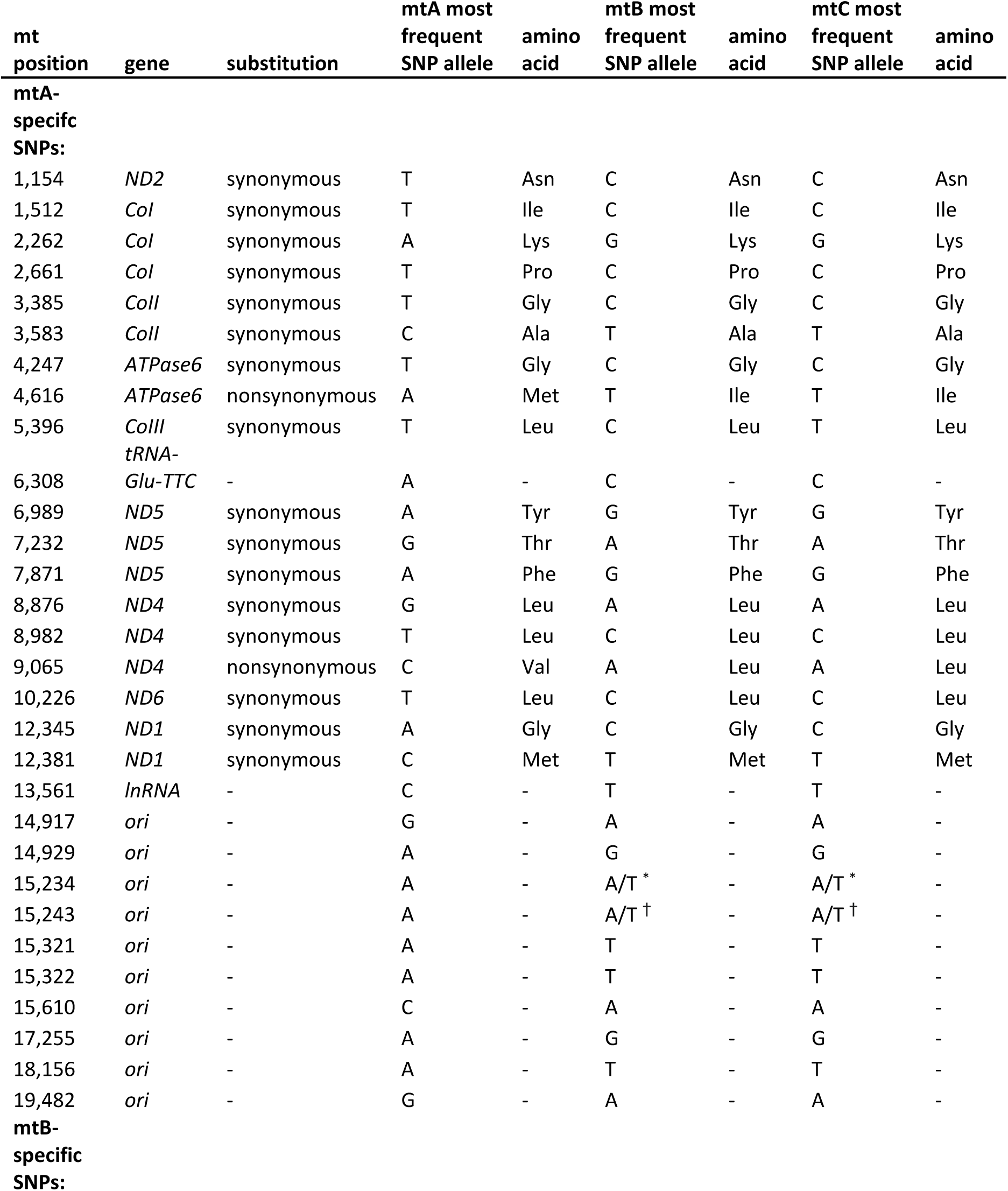
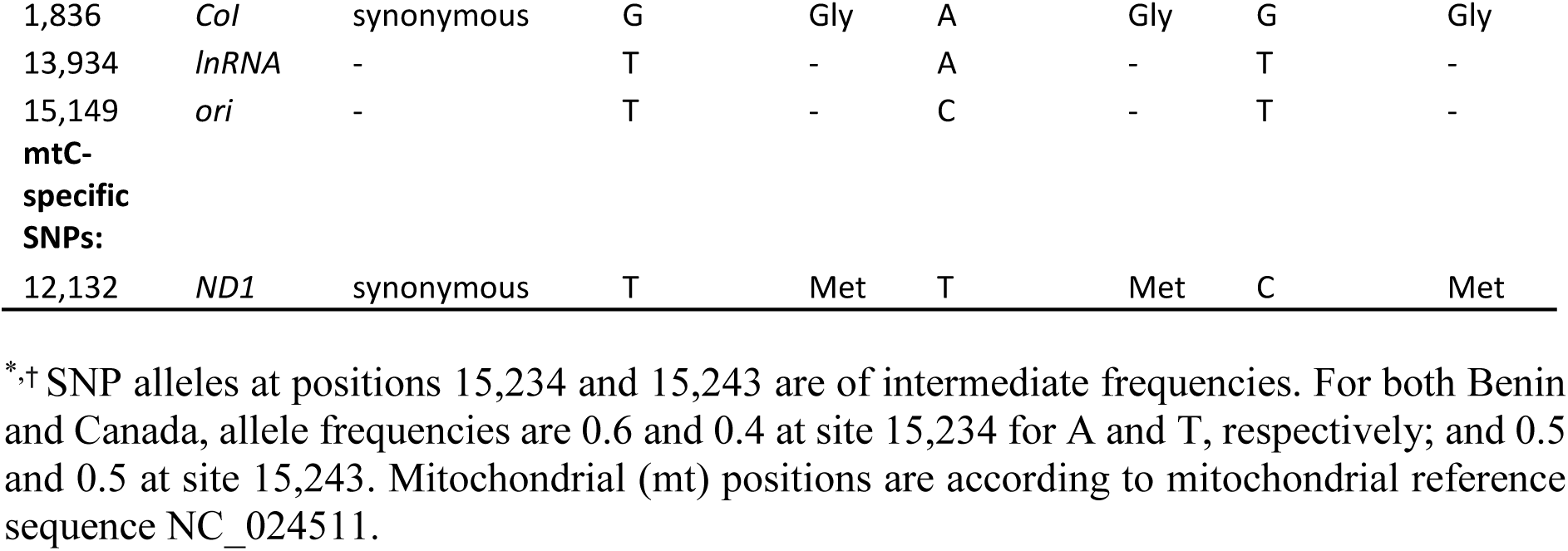
Annotation of significantly differentiated mitochondrial SNPs (Fisher’s exact test; FDR=0.1%) derived from each population (Australia:A, Benin:B, Canada:C). Population replicates carrying outlier mitotypes AA3, CA1 and CA3 were excluded. Mitochondrial (*mt*) *position* indicates the location of the SNP within the mitochondrial genome according to mitochondrial reference sequence NC_024511. *Gene* indicates the name of the gene the SNP is located in. *Substitution* indicates whether the nucleotide change results in a change in the amino acid encoded; with synonymous indicating no change, and non-synonymous indicating a change. *Most frequent SNP allele* indicates the most frequent nucleotide present for a given positional location for mtDNA captured from each population. *Amino acid* indicates the identity of the amino acid encoded by a particular SNP.

The cyto-nuclear populations harbouring Australian cytoplasms possessed the largest amount of mitochondrial genotypic variation and differentiation, relative the other cyto-nuclear populations harbouring cytoplasms from the other two populations (Fig. 1, Fig. 2, Supporting Information Fig. 3-5). The mitochondrial genetic variation within populations harbouring Beninese and Canadian cytoplasms were relatively similar to each other, with only three significantly differentiated sites separating the populations (Fig. 2). The associated genotype network, based on the 34 SNPs and the consensus sequences of the 27 populations, reveals most of the populations cluster according to the location of origin of their cytoplasm (Fig. 1). Exceptions are population replicates CA1 and CA3 whose mitochondrial sequences appeared more closely related to cyto-nuclear populations harbouring the Benin cytoplasm, rather than the Canadian type (Supporting Information Fig. 3-5).

**Figure 2.**
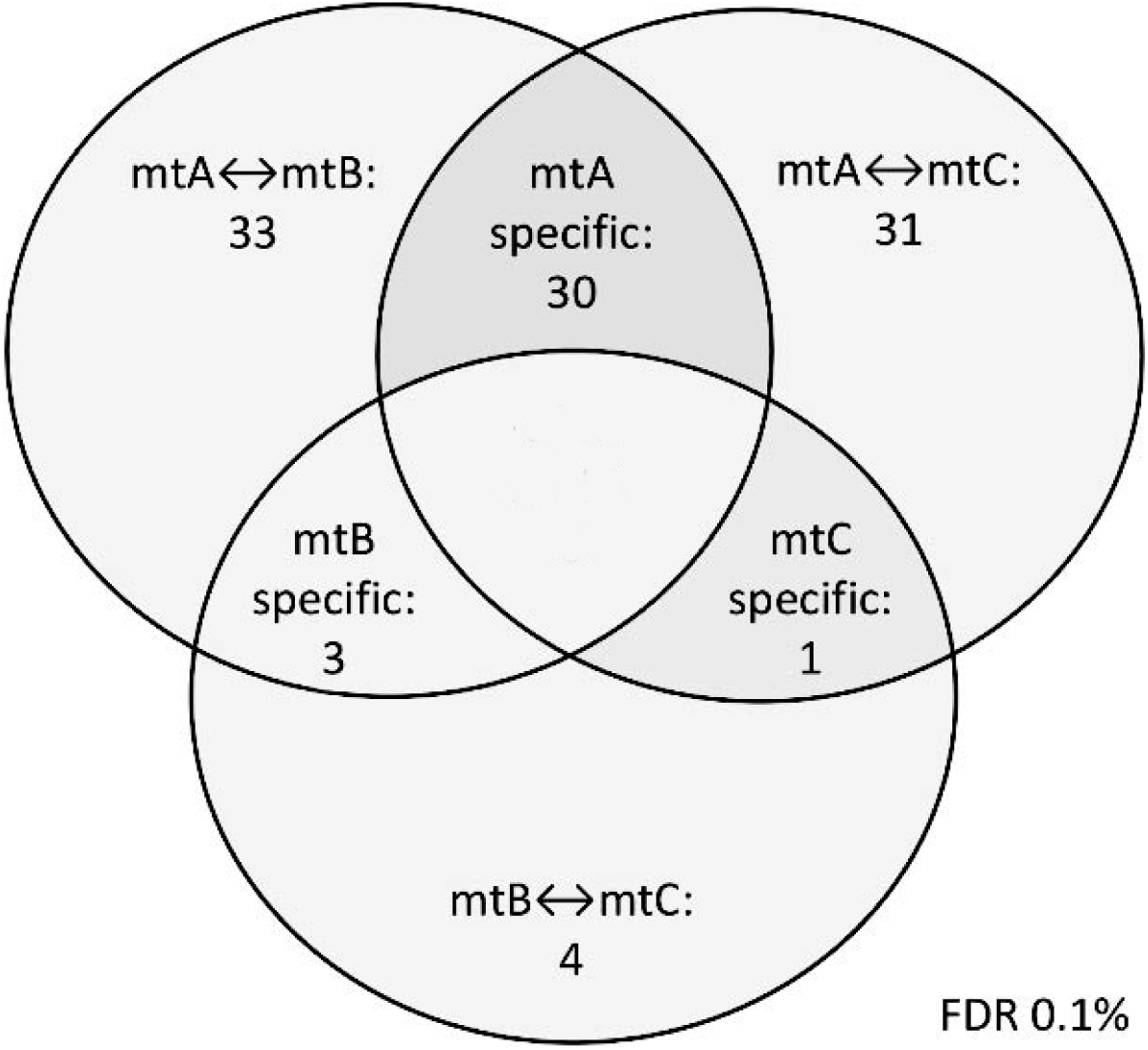
Diagram showing the numbers of significantly differentiated SNPs between mitochondrial genomes from Australia, Benin, and Canada. Numbers of variants specific to either Australia, Benin or Canada are given as well. Those variants are in high frequency in lines containing mtDNA derived from one origin, but are absent or in low frequency in lines with mtDNA from the other two origins.

Furthermore, two distinct mitotypes were present in the cyto-nuclear populations harbouring Australian cytoplasms. These same mitotypes have previously been characterized as distinct haplotypes (Camus et al., 2017; Lajbner et al., 2018), with one of the haplotypes, denoted A1, found to be predominate in low latitude sub-tropical regions of Australia and another haplotype denoted B1, present in higher latitude temperate regions of Australia (Camus et al., 2017). The B1 haplotype was nearly fixed (in very high frequency) in eight out of the nine cyto-nuclear populations carrying Australian cytoplasms, with the A1 haplotype nearly fixed in one population replicate (AA3, Supporting Information Fig. 3).

### Partitioning of cytoplasmic and nuclear sources of variance

Lifespan was affected by a significant interaction between the cytoplasm, nuclear background and sex of the individuals (Table 3A, Cyto × Nuclear × Sex: *Χ*^2^_4_ = 11.150*, P* = 0.0249). The combinations of cyto-nuclear genotype that encoded longest life in females were not the same combinations that encoded longest life in males (Fig 3., Fig. 4, Supporting Information Fig. 6).

**Figure 3.**
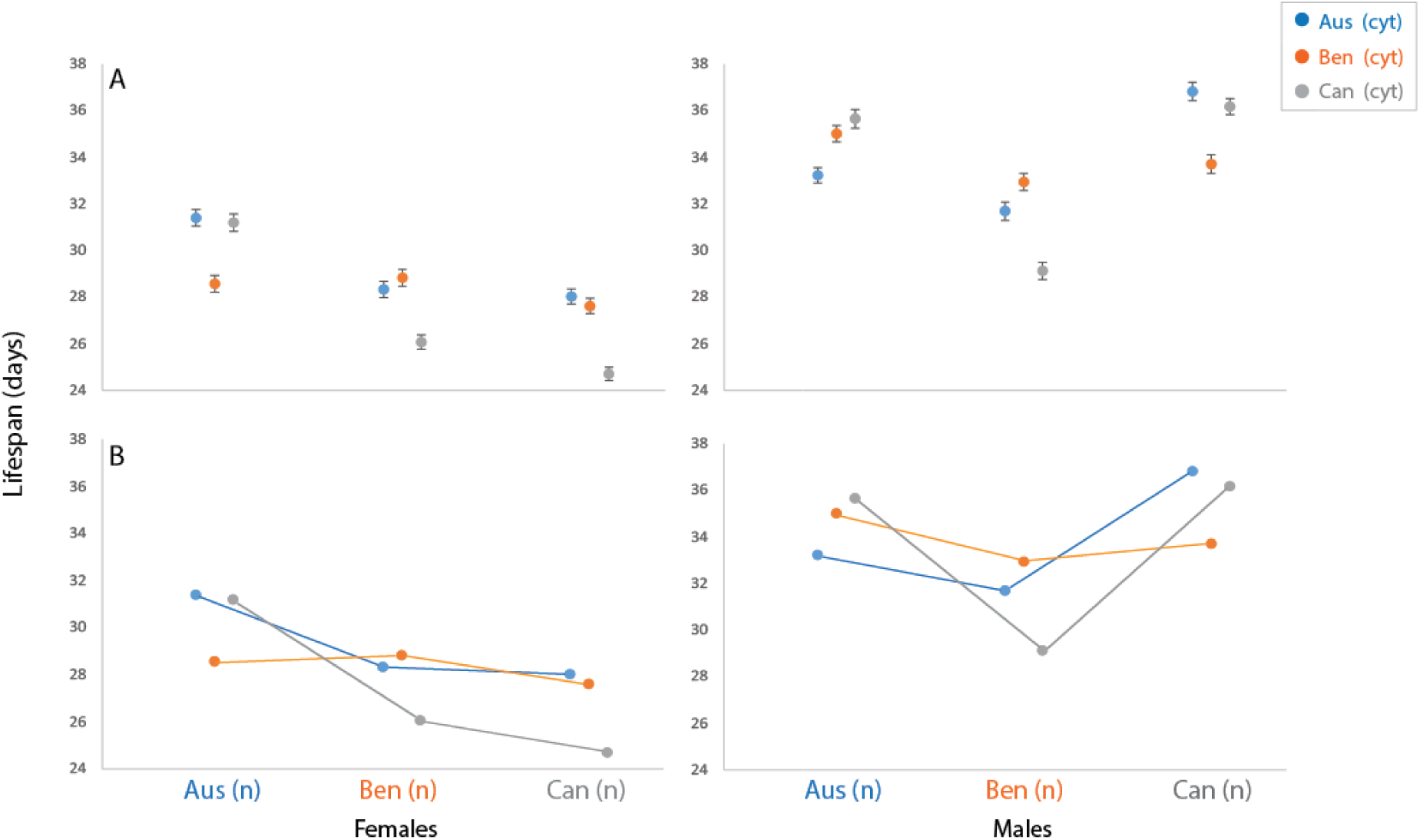
Cyto-nuclear effects on longevity. **A**. Mean lifespan (± 1 standard error) of each of the three cytoplasms expressed against each of the three nuclear backgrounds, in females (left hand panel) and males (right hand panel). Lifespan (as measured in days) is indicated on the vertical axis, with the nuclear background denoted on the horizontal axis. Each colour denotes a different cytoplasm: Australia shown in blue, Benin shown in orange, Canada shown in grey. **B**. Reaction norms of the same data. Mean lifespan is indicated on the vertical axis, and nuclear background on the horizontal axis.

**Figure 4.**
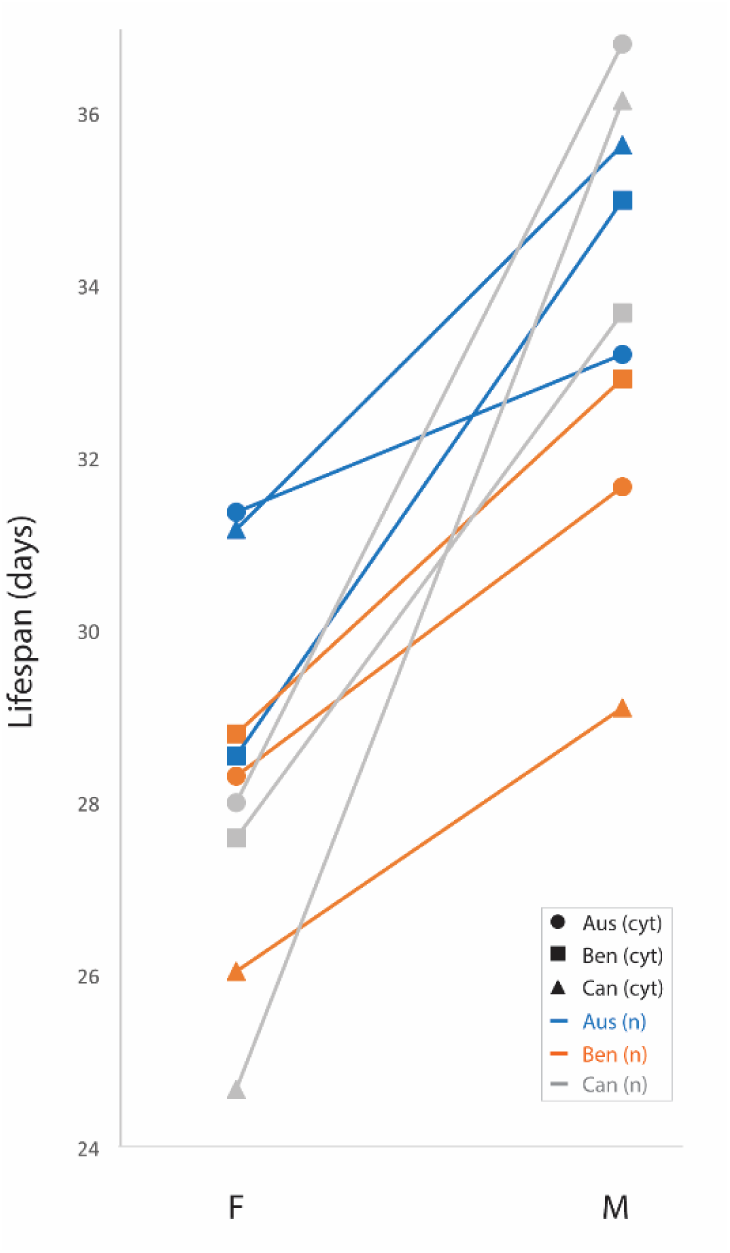
Performance of cyto-nuclear genotypes for the two sexes. Interaction plot of mean lifespan per sex across all cyto-nuclear genotypes. The nuclear genetic background originating from Australia is shown in blue, the nuclear genetic background originating from Benin is shown in orange, the nuclear genetic background originating from Canada is shown in grey. Cytoplasms originating from Australia shown with circles, cytoplasms originating from Benin shown with shown with squares, cytoplasms originating from Canada shown with triangles. Sexes, indicated by F or M.

For example, relative to the other two cytoplasms the Beninese cytoplasm conferred short lifespan in females when expressed in the Australian nuclear background, but high relative male lifespan in this same nuclear background. The Canadian cytoplasm encoded markedly shorter lifespan in females when expressed in its matched Canadian nuclear background, but high lifespan in males, relative to other cyto-nuclear combinations. Thus, reaction norms of cytoplasm performance were discordant across sexes within the Australian and Canadian nuclear backgrounds. In contrast, the reaction norms converged within the Beninese nuclear background; the Canadian cytoplasm conferred the shortest lifespan in both of the sexes in this background (Fig. 3, Supporting Information Fig. 6).

Males generally outlived females under the mixed-sex conditions in which the individuals were maintained over the course of the experiment. Notably, however, the sex differences in lifespan depended on the specific cyto-nuclear combination; in some cases sex differences were eroded, such as the case of individuals harbouring Australian cytoplasms in the Australian nuclear background (Fig. 3A).

### Effects of cyto-nuclear mismatching

We found no evidence that evolutionary matched (putatively ‘coevolved’) combinations of cyto-nuclear genotype conferred longer life than evolutionary mismatched (‘mismatched’) combinations, in either sex (Table 3B, Coevolutionary status: *X*^2^_1_ = 0.0315, *P* > 0.05, interaction between coevolutionary status and sex was dropped from the final model: *X*^2^_1_ = 0.0967*, P* = 0.756).

### Partitioning mitochondrial from *Wolbachia* effects

The screen for *Wolbachia* revealed that all of the Australian cytoplasms are infected while the others harnessed here were not. Furthermore, genotyping of the strains revealed that two of the population replicates of the Canadian cytoplasm with the Australian nuclear background were fixed for alternative mitotypes compared to the other strains carrying Canadian cytoplasms. Given this, a key consideration then becomes discerning whether the cytoplasmic effects observed were primarily attributable mitochondrial sequence variation or *Wolbachia* infection. Indeed, one of the *Wolbachia*-infected population replicates (AA3: Australian cytoplasm in Australian nuclear background, replicate 3) carries an alternative haplotype (A1) than the other *Wolbachia*-infected population replicates, and this provided an opportunity to explore whether lifespan associated with this population differed from that of the other populations carrying Australian cytoplasms within Australian nuclear background (AA1 and AA2, which harbor the B1 haplotype), and also to compare the effects of the Australian cytoplasms (which are all infected with *Wolbachia*) to those of the Canadian and Benin mitotypes (none of which are infected with *Wolbachia*, namely BA1-3, CA1&3, CA2) within the same Australian nuclear background.

The interaction between mitotype and sex of the individuals was not statistically significant, although the *P* value approached the 0.05 alpha criterion (Table 4, Mito × Sex: *X*^2^_4_ = 9.4313, *P* = 0.0512), and a general effect of mitotype was not found to be significant. However, visual inspection of the pattern of effects across the haplotypes (Fig. 5) suggests that the contrast between the mean lifespan associated with the *Wolbachia*-infected Australian cytoplams – i.e. AA3 (which harbours the A1 haplotype) and AA1&AA2 (which harbours the B1 haplotype) – would suggest that at least some of the cytoplasmic-mediated effects described above may be directly encoded by sequence differences between mitotypes.

**Figure 5.**
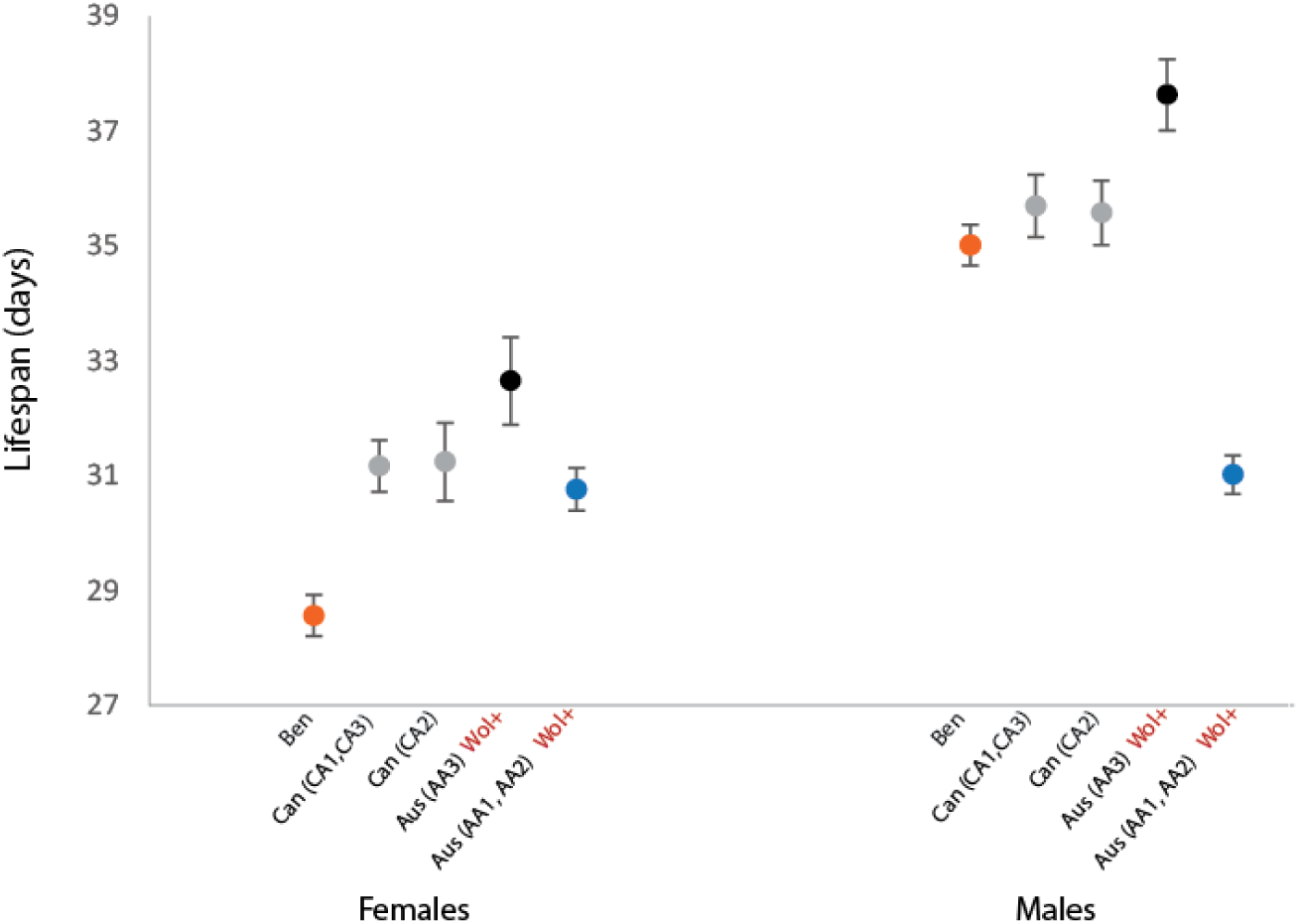
Mean lifespan of mitotypes when expressed alongside the Australian nuclear background, in females and males. Comparisons are: 1) Beninese (comprising BA1, BA2, BA3), 2) Canadian CA1 and CA3 (whose prevailing mitotypes were more similar to B mitotypes than to the other C mitotypes 3) Canadian CA2, 4) Australian A1 (AA3), and 5) B1 (AA1 and AA2). Beninese cytoplasms shown in orange, Canadian shown in grey, Australian lines carrying the A1 haplotype shown in black, Australian lines carrying the B1 haplotype shown in blue. Wol+ in red lettering indicates *Wolbachia* infected lines, all others are free from *Wolbachia* infection. Error bars indicate standard error based on calculated means.

**Table 3.**
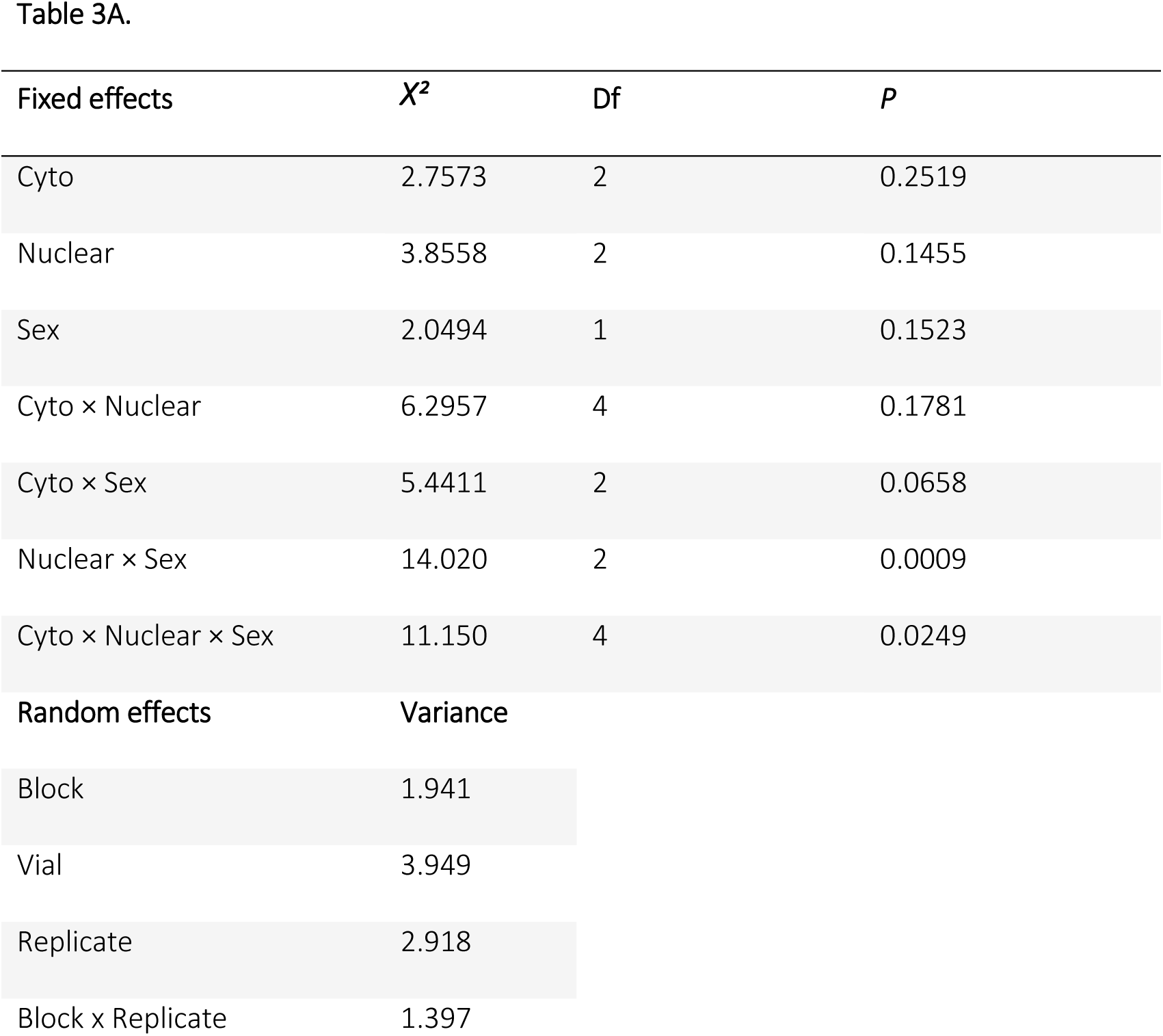
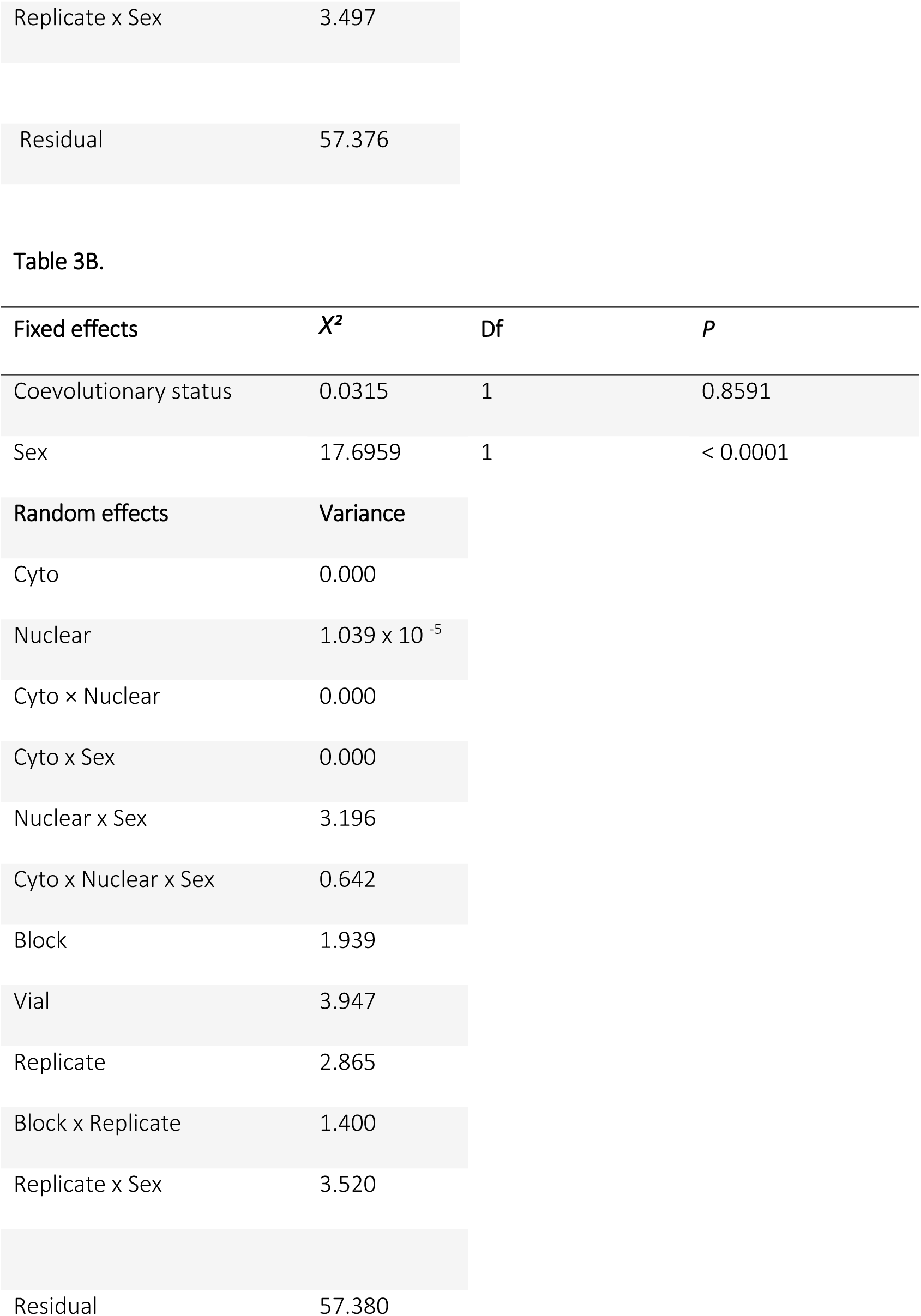
A. Effects of genotype and sex on lifespan across the 27 cytonuclear population replicates. Linear mixed effects model of lifespan (measured in days) with fixed effects of cytoplasm (Cyto), nuclear background (Nuclear), sex (Sex) and interactions between the three. Interactions included those between cytoplasm and nuclear background (Cyto × Nuclear) cytoplasm and sex (Cyto × Sex), nuclear background and sex (Nuclear × Sex), and three-way interactions between cytoplasm, nuclear background and sex (Cyto × Nuclear × Sex). Random effects were experimental block (Block), vial identity (Vial), cyto-nuclear population replicate (Replicate), the interaction between experimental block and cyto-nuclear population replicate (Block × Replicate), and the interaction between cyto-nuclear population replicate and sex (Replicate × Sex). **B**. Effects of experimental mismatching of cyto-nuclear genotypes on longevity. Fixed effects are coevolutionary status (Status) (coevolved or mismatched) and sex. Random effects included cytoplasm (Cyto) and nuclear background (Nuclear), any interactions between the the two and also all possible interactions with sex (Sex). Interactions of random effects included those between cytoplasm and nuclear background (Cyto × Nuclear), cytoplasm and sex (Cyto × Sex), nuclear background and sex (Nuclear × Sex) and three-way interactions between cytoplasm, nuclear background and sex (Cyto × Nuclear × Sex). Other random effects included block (Block), vial identity (Vial), cyto-nuclear population replicate (Replicate), the interaction between experimental block and cyto-nuclear population replicate (Block × Replicate), and the interaction between cyto-nuclear population replicate and sex (Replicate × Sex).

| Fixed effects | $X^2$ | Df | $P$ |
| --- | --- | --- | --- |
| Cyto | 2.7573 | 2 | 0.2519 |
| Nuclear | 3.8558 | 2 | 0.1455 |
| Sex | 2.0494 | 1 | 0.1523 |
| Cyto × Nuclear | 6.2957 | 4 | 0.1781 |
| Cyto × Sex | 5.4411 | 2 | 0.0658 |
| Nuclear × Sex | 14.020 | 2 | 0.0009 |
| Cyto × Nuclear × Sex | 11.150 | 4 | 0.0249 |
| **Random effects** | **Variance** |  |  |
| Block | 1.941 |  |  |
| Vial | 3.949 |  |  |
| Replicate | 2.918 |  |  |
| Block x Replicate | 1.397 |  |  |

|  |  |
| --- | --- |
| Replicate x Sex | 3.497 |
| Residual | 57.376 |

| **Fixed effects** | **$X^2$** | **Df** | ***P*** |
| --- | --- | --- | --- |
| Coevolutionary status | 0.0315 | 1 | 0.8591 |
| Sex | 17.6959 | 1 | < 0.0001 |

| **Random effects** | **Variance** |
| --- | --- |
| Cyto | 0.000 |
| Nuclear | $1.039 \times 10^{-5}$ |
| Cyto x Nuclear | 0.000 |
| Cyto x Sex | 0.000 |
| Nuclear x Sex | 3.196 |
| Cyto x Nuclear x Sex | 0.642 |
| Block | 1.939 |
| Vial | 3.947 |
| Replicate | 2.865 |
| Block x Replicate | 1.400 |
| Replicate x Sex | 3.520 |
| Residual | 57.380 |

**Table 4.** Effects of mitotype and sex on lifespan for the analysis of the Australian nuclear background only. Linear mixed effects model of longevity (measured in days) with fixed effects of mitotype identity (Mito), sex (Sex) and the interaction between mitotype and sex (Mito × Sex). Random effects were experimental block (Block), vial identity (Vial), population replicate (Replicate), the interaction between experimental block and population replicate (Block × Replicate), the interaction between population replicate and sex (Replicate × Sex), the interaction between mitotype, sex and experimental block (Mito × Nuclear × Sex), the interaction between mitotype and experimental block (Mito × Block), and the interaction between sex and experimental block (Sex × Block).

| Fixed effects | $X^2$ | Df | $P$ |
| --- | --- | --- | --- |
| Mito | 2.8023 | 4 | 0.5914 |
| Sex | 0.0325 | 1 | 0.8569 |
| Mito x Sex | 9.4313 | 4 | 0.0512 |
| **Random effects** | **Variance** |  |  |
| Block | 1.945 |  |  |
| Vial | 1.980 |  |  |
| Replicate | 6.045 |  |  |
| Block x Replicate | 1.739 |  |  |
| Replicate x Sex | 4.047 |  |  |
| Mito x Sex x Block | 1.041 |  |  |
| Mito x Block | 0.000 |  |  |
| Sex x Block | 0.000 |  |  |
| Residual | 61.946 |  |  |

## DISCUSSION

The interactions between the mitochondrial and nuclear genome are key drivers of critical metabolic processes in eukaryotes (Bar-Yaacov et al., 2012; Levin et al., 2014; Wolff et al., 2014; Stojković & Đordević, 2017; Mossman et al., 2019; St. John, 2019), and have been increasingly invoked as mediators of fundamental evolutionary processes ranging from the evolution of reproductive isolation and speciation (Gershoni et al., 2009; Chou & Leu, 2010; Burton & Barreto, 2012; Burton et al., 2013; Hill, 2016; Hill, 2017; Haddad et al., 2019; Lima et al., 2019; Tobler et al., 2019), to adaptation under sexual selection (Hill & Johnson, 2013; Hill, 2015), to the evolution of sex differences (Camus et al., 2012; Vaught & Dowling, 2018; Montooth et al., 2019), and evolution of sexual reproduction (Havird et al., 2015; Radzvilavicius & Blackstone, 2015; Radzvilavicius, 2016). A number of studies has previously reported evidence that disruption of putatively coevolved cytoplasmic and nuclear genomes leads to negative effects on phenotypic expression in eukaryotes ranging from yeast to humans (Hill et al., 2019). However, for those studies conducted in metazoans to date, many have focused on the consequences of cyto-nuclear mismatching between highly divergent genetic lineages, while others have focused on only one of the two sexes. Accordingly, it remains unclear whether the magnitude and patterns of cyto-nuclear interactions may manifest differently in each of the two sexes within the same species.

The goal of this study was therefore to test whether cyto-nuclear interactions affected the expression of lifespan across three populations of *D. melanogaster*, and to explore whether the direction or magnitude of such interactions was sex-specific. Consistent with previous reports in fruit flies, copepods, other invertebrates, and mice (reviewed in Dowling et al., 2008; Burton et al., 2013; Dowling, 2014; Wolff et al., 2014; Hill, 2015; Stojković & Đordević, 2017; Hill et al., 2019), we found that phenotypic expression of a key component of life-history, lifespan, was contingent on epistatic interactions between cytoplasmic and nuclear genes. This result is consistent with previous studies to have reported cyto-nuclear interactions for lifespan and survival in fruit flies such as *D. melanogaster* and *D. simulans* (James & Ballard, 2003; Rand et al., 2006; Clancy et al., 2008; Zhu et al., 2014; Jelić et al., 2015; Drummond et al., 2019), seed beetles (Dowling et al., 2010; Đorđević et al., 2015, 2017), and nematodes (Zhu et al., 2015). Many of these studies, however, either examined lifespan effects in only one sex or did not incorporate a statistical approach that enabled a direct comparison of the patterns and magnitude of cyto-nuclear epistasis across the sexes. Here, we found that cyto-nuclear interactions contribute to the expression of lifespan and that the outcomes of these interactions differ across the sexes. Our results extend on a few recent studies that have observed sex-dependent mito-nuclear effects on longevity. Firstly, Drummond *et al.,* (2019) reported that the effects of mtDNA haplotype on survival were mediated by the nuclear background in *D. melanogaster*, with evidence that the magnitude of the mito-nuclear interactions was greater in males than females. Đorđević *et al*., (2015, 2017) found these same mito-nuclear and sex-specific effects for lifespan in genetic lines of seed beetles carrying combinations of mitochondrial and nuclear genomes selected for either long or short life. Finally, Jelić *et al.,* (2015) reported similar patterns present in *Drosophila subobscura*, with certain sympatric mito-nuclear combinations conferring high or low lifespan in one sex but not the other. In our study we found that, no cytoplasm conferred generally superior lifespan across nuclear backgrounds; the combinations of cyto-nuclear genotype that conferred longest life in males were not the same combinations conferring longest life in females. Furthermore, we observed no sex-specificity in the magnitude of cyto-nuclear effect size for longevity.

We attempted to address whether the cyto-nuclear interactions observed in our study were mediated primarily by heterogeneity in the *Wolbachia* infection status of the populations, since *Wolbachia* has been observed to positively or negatively influence lifespan outcomes across previous studies (Fry & Rand, 2002; Fry et al., 2004; Alexandrov et al., 2007; Capobianco et al., 2018), or by sequence divergence in the mitochondrial genome, across the cyto-nuclear populations. To this end, we were able to address two questions. First, whether or not the major differences in lifespan across populations grouped in two clusters of cyto-nuclear populations – those that carry *Wolbachia* infection (those with cytoplasms of Australian origin) and those that do not (all other populations). We observed that populations with the Australian cytoplasm do not behave consistently differently relative to populations with Beninese or Canadian cytoplasmic backgrounds, suggesting that *Wolbachia* is not having a disproportionately large or consistent role in their effects on lifespan. Secondly, within each cluster (*Wolbachia*-infected versus uninfected groups) of populations, we were able to examine whether differences in phenotypic expression across the cyto-nuclear populations mapped to the prevailing mitochondrial haplotype of each population. Specifically, we focused on the set of cyto-nuclear populations expressed in the Australian nuclear background. In this nuclear background, we found that one *Wolbachia*-infected cytoplasm (AA3) carried a different prevailing mitochondrial haplotype [a haplotype previously denoted A1 by Camus et al., (2017)] than the other infected cytoplasms (AA1 and AA2) that carried the B1 haplotype. The overall mean of the population carrying the B1 haplotype was putatively lower than those carrying the A1 haplotypes, and this effect seemed amplified in males (Hedges gmales = 0.92 ± 0.18, Hedges g_females_= 0.22 ± 0.18), albeit the interaction between haplotype and sex was not statistically significant (*P* = 0.053). The genotyping analysis indicates the B1 haplotype carries 20 unique SNPs that distinguish it from the Benin and Canada mitotypes (Fig. 2). Interestingly, some of these unique changes occurred within the origin of replication as well as in the tRNA^Glu^ (Fig. 1). These genetic differences might underlie the phenotypic differences we observed, however it’s difficult to confirm such a possibility in the present study.

We predicted that any cyto-nuclear incompatibilities that manifest upon experimental disruption of coevolved cyto-nuclear genotypes might exhibit signatures of male-bias. This prediction is based on the hypothesis that maternal inheritance of the mitochondrial genome will facilitate the accumulation of mutations that are male-harming, but benign or beneficial to females (Frank & Hurst, 1996; Gemmell et al., 2004; Beekman et al., 2014; Dowling & Adrian, 2019); an evolutionary prediction that has received some empirical support from recent studies in *D. melanogaster* (Camus et al., 2012; Innocenti et al., 2011; Patel et al., 2016; Camus & Dowling, 2018; Drummond et al., 2019; Nagarajan-Radha et al., 2019 in-press), hares (Smith et al., 2010), and humans (Martikainen et al., 2017; Milot et al., 2017; Gudiseva et al., 2019). We found no consistent evidence that mismatching of population-specific cytoplasmic and nuclear genotypes results in a decrease in lifespan in either of the sexes. This does not exclude the possibility that putatively mismatched cyto-nuclear combinations could have conferred negative effects on other key traits not assessed in this study, such as reproduction, metabolic rate or ROS production. However, the results are consistent with the idea that the levels of mitochondrial genetic divergence observed within many species may not be sufficient to drive mito-nuclear mediated reproductive isolation between populations, a plausible outcome in the face of gene flow between these populations. Such a pattern would be congruent with a growing body of similar studies conducted at the intra-specific scale, in *Drosophila* and *Callosobruchus* beetles, which have reported that although cyto-nuclear interactions are present, the experimental mismatching of putatively coevolved combinations does not consistently lead to poorer phenotypic outcomes (Dowling et al., 2007; Arnqvist et al., 2010; Jelić et al., 2015). Notwithstanding, some studies examining the consequences of mito-nuclear mismatching at the intraspecific scale have documented clear evidence for fitness impairment in some species; albeit these studies exhibit much higher levels of mtDNA sequence divergence than seen between populations of humans or *D. melanogaster* (Morrow et al., 2015). This includes studies of the intertidal copepod *T. californicus*, which exhibits mtDNA sequence divergence of up to 20% between populations, and the nematode *C. briggsae*, which exhibits divergence of around 2.6% for protein-coding genes (Chang et al., 2016; Haddad et al., 2018; Adineh & Ross, 2019; Cazares-Navarro & Ross, 2019). Currently, however, it remains unclear whether increases in levels of mtDNA sequence divergence between populations are likely to routinely drive the emergence of mito-nuclear incompatibilities; since several studies comparing combinations of mito-nuclear genotype sourced from different species of *Drosophila* (*D. simulans* and *D. melanogaster*, whose mtDNA sequences are ∼5% divergent (Solignac et al., 2004)), did not detect fitness reductions in flies carrying interspecific combinations (Zhu et al., 2014; Mossman et al., 2016a, b; Mossman et al., 2019). As such, the likelihood of mito-nuclear incompatibility occurring, or the degree of mito-nuclear incompatibility, may not necessarily correspond to the amount of genetic divergence present, and more work needs to resolve these issues within and across species (Havird et al., 2019; Hill et al., 2019).

In conclusion, our study provides some new insights into the dynamics and evolutionary implications of cyto-nuclear interactions, by showing cyto-nuclear epistasis contributes to lifespan even at the intraspecies scale, and that the magnitude and pattern of effects across cyto-nuclear combination can be sex-specific. This suggests that selection for optimally performing cyto-nuclear genotypes will in part be sex-specific, which is likely to be the case given the mitochondrial genome is transmitted only through females, and can, therefore, only respond to selection directly through females. Currently, several questions remain. It is unclear how much of the cyto-nuclear effect on life-history trait expression is explicitly tied to mitochondrial sequence divergence; or how much mitochondrial divergence is required to precipitate fitness interactions with the nuclear genotype. Recently, it has even be argued that cyto-nuclear interactions will be relevant to molecular processes that are at play within the individual (Havird et al., 2019; Shtolz & Mishmar, 2019; Wei et al., 2019), even occurring via interactions between gene products encoded by different mtDNA molecules, present in heteroplasmy within the cell, and nuclear-encoded products (Wolff et al., 2014), and to epigenetic mechanisms (Grunau et al., 2018; Kopinski et al., 2019). Ultimately, the degree to which such interactions may contribute to the missing variance in quantitative genetic studies remains to be fully investigated (Rand et al., 2018; Zhu et al., 2019). Finally, it is now clear that several extra-nuclear sources of variance may affect the expression of the organismal phenotype, and it is possible that these cytoplasmic genetic elements interact not only with genetic variants in the nuclear background to shape trait values via cyto-nuclear interactions, but also epistatically with each other, which could manifest in complex cytoplasmic-by-cytoplasmic-by-nuclear interactions underpinning variation in organismal function (Labjner et al., 2018; Havird et al., 2019). Such possibilities deserve experimental attention.

## Acknowledgements

We are grateful to members of the Dowling and Sgró labs for their support in the lab work associated with this project, and four anonymous reviewers for comments on a previous version of the manuscript.

