## Supporting Information for "Interactions between cytoplasmic and nuclear genomes confer sex-specific effects on lifespan in *Drosophila melanogaster*"

### MATERIALS AND METHODS

#### Partitioning of cytoplasmic and nuclear sources of variance

Survival data for the 27 cyto-nuclear population replicates was analysed using the *coxme* package (Therneau, 2018) in R. A *surv* model (Therneau, 2015) was constructed with the same designations of fixed and random effects as described in the methods for lifespan. The model included fixed effects of cytoplasmic origin, nuclear background, and sex, as well as interactions between the three. Random effects, describing the hierarchical structure of the data, included experimental sampling block (n = 6), the vial identity in which the assays took place (n = 645), and the cyto-nuclear population replicate (n = 27). Interactions involving random effects were not included since the *coxme* package is unable to model such interactions, therefore the usual model simplification process did not apply to this analysis for the random effects statement. Fixed effects in the final model were estimated using a Wald's Type III sums-of-squares analysis and random effects were estimated using the *summary* function.

In order to visualize the survival outcomes of the three cytoplasms (Australia, Benin and Canada) within each of the three nuclear backgrounds (Australia, Benin and Canada) for males and females, Kaplan-Meier survival curves were generated using a protocol adapted from (https://github.com/michaelway/ggkm/), by first fitting a basic survival model for the time-to-death in days using the *surv* function within the *Survival* package (Therneau, 2015) and then plotting the fitted data with the *ggkm* function within *ggplot2* (Wickham, 2016).

#### Homing in on mitochondrial genetic effects

The results of the genotyping analysis revealed that one of cyto-nuclear population replicates harbouring Australian cytoplasms were nearly fixed for a different mitotype than the others (A1 haplotype in AA3, B1 haplotype in the others), and two of the cyto-nuclear populations with Canadian cytoplasms contained mitotypes that were more similar to those of Benin than Canada. Two potential analytical approaches could be employed to attempt to further investigate the cytoplasmic sources of variance for lifespan, to determine whether patterns of lifespan could be mapped to the level of the mitochondrial genotype. One approach would to be to exclude the population replicates harbouring three 'outlier' mitotypes (CA1 and CA3, which appear to be more closely related to Benin than Canada; and AA3, which harbors the A1 haplotype) from the analyses, while the other approach would be to exclude AA3 (that has the Australian A1 haplotype) but recode the cytoplasmic contribution of the CA1 and CA3 to that of Beninese origin, given the similarities between the mitochondrial profiles of CA1 and CA3 with the populations harbouring Beninese cytoplasms. We chose to exclude the outlier population replicates (AA3, CA1 and CA3) from this second analysis (thus, this analysis is based on 24, rather than 27 replicates), but we note that the alternative approach (of recoding CA1 and CA3) led to the same qualitative conclusions.

### RESULTS

#### Partitioning of cytoplasmic and nuclear sources of variance

The results of the survival analyses largely mirrored the results uncovered for lifespan (Supporting Information Table 1, Supporting Information Fig. 7-12) and furthermore this survival analysis did not reveal the presence of any effects that could have potentially confounded the effects uncovered for lifespan (Supporting Information Fig. 7-12). Analogously to lifespan, survival was found to be affected by an interaction between the cytoplasm, nuclear background and sex of the individuals (Supporting Information Table 1, Cyto × Nuclear × Sex: *Wald* = 42.8540, $P < 0.0001$) and the sex-specific survival outcomes of the cyto-nuclear combinations matched those of mean lifespan outcomes (Supporting Information Fig. 7-12).

**Homing in on mitochondrial genetic effects**

The same patterns uncovered for the global analysis of the dataset including all 27 cyto-nuclear replicate population were upheld when reanalysing the dataset, having excluded AA1, CA2 and CA3 populations whose mitotypes were discordant with the other cyto-nuclear populations harboring respective Australian and Canadian cytoplasms. For this analysis where the population replicates harbouring three ‘outlier’ mitotypes were excluded, the overall 3-way interaction of cytoplasm, nuclear background and sex of the individual was significant (Supporting Information Table 2A, Cyto × Nuclear × Sex: $X^2_4$ = 15.8968, $P$ = 0.0032; Supporting Information Fig. 2).

### SUPPORTING TABLES AND FIGURES

**Supporting Information Table 1.** Partitioning of cytoplasmic and nuclear sources of variance. Effects of genotype and sex on survival. Survival data were analysed using Cox mixed effects models of survival with fixed effects of cytotype identity (Cyto), nuclear background (Nuclear), sex (Sex) and interactions between the three. Interactions included those between cytotype and nuclear background (Cyto × Nuclear), cytotype and sex (Cyto × Sex), nuclear background and sex (Nuclear × Sex), and three-way interactions between cytotype, nuclear background and sex (Cyto × Nuclear × Sex). Random effects were experimental block (Block), vial identity (Vial), and cyto-nuclear population replicate (Replicate).

SI Table 1.

| Fixed effects | Wald | Df | *P* |
|---|---|---|---|
| Cyto | 3.4167 | 2 | 0.1812 |
| Nuclear | 8.1844 | 2 | 0.0167 |
| Sex | 2.4554 | 1 | 0.1171 |
| Cyto × Nuclear | 9.7697 | 4 | 0.0445 |
| Cyto × Sex | 22.7656 | 2 | < 0.0001 |
| Nuclear × Sex | 61.5288 | 2 | < 0.0001 |
| Cyto × Nuclear × Sex | 42.8540 | 4 | < 0.0001 |
| **Random effects** | **Variance** | | |
| Block | 0.033 | | |
| Vial | 0.141 | | |
| Replicate | 0.070 | | |

**Supporting Information Table 2.** Homing in on mitochondrial genetic effects. **A**. Effects of genotype and sex on lifespan when the three outlier populations (AA3, CA1 and CA3) were omitted. Linear mixed effects model of longevity (measured in days) with fixed effects of cytotype identity (Cyto), nuclear background (Nuclear), sex and interactions between the three. Interactions included those between cytotype and nuclear background (Cyto × Nuclear), cytotype and sex (Cyto × Sex), nuclear background and sex (Nuclear × Sex), and three-way interactions between cytotype, nuclear background and sex (Cyto × Nuclear × Sex). Random effects were experimental block (Block), vial identity (Vial), cyto-nuclear population replicate (Replicate), the interaction between experimental block and cyto-nuclear population replicate (Block × Replicate), and the interaction between cyto-nuclear population replicate and sex (Replicate × Sex). **B**. Effects of mito-nuclear status on lifespan when the three outlier populations (AA3, CA1 and CA3) were omitted. Linear mixed effects model of longevity (measured in days) with fixed effects of coevolutionary status (Status) (coevolved or mismatched) and sex (Sex). Random effects included cytotype identity (Cyto) and identity of the nuclear background (Nuclear), any interactions between the the two and also all possible interactions with sex.

SI Table 2A.

| Fixed effects | $X^2$ | Df | *P* |
|---|---|---|---|
| Cyto | 2.2978 | 2 | 0.3169 |
| Nuclear | 2.7126 | 2 | 0.2576 |
| Sex | 0.0351 | 1 | 0.8514 |
| Cyto × Nuclear | 5.7680 | 4 | 0.2172 |
| Cyto × Sex | 9.6097 | 2 | 0.0082 |
| Nuclear × Sex | 20.0216 | 2 | < 0.0001 |
| Cyto × Nuclear × Sex | 15.8968 | 4 | 0.0032 |
| **Random effects** | **Variance** | | |
| Block | 1.691 | | |
| Vial | 3.997 | | |
| Replicate | 1.570 | | |
| Block x Replicate | 1.366 | | |
| Replicate x Sex | 3.102 | | |
| Residual | 55.727 | | |

SI Table 2B.

| Fixed effects | $X^2$ | Df | *P* |
|---|---|---|---|
| Coevolutionary status | 0.0025 | 1 | 0.9601 |
| Sex | 17.9823 | 1 | < 0.0001 |
| **Random effects** | **Variance** | | |
| Cyto | $1.573 \times 10^{-13}$ | | |
| Nuclear | 0.000 | | |
| Cyto × Nuclear | $4.836 \times 10^{-2}$ | | |
| Cyto x Sex | $5.813 \times 10^{-12}$ | | |
| Nuclear x Sex | 2.509 | | |
| Cyto x Nuclear x Sex | 1.471 | | |
| Block | 1.688 | | |
| Vial | 3.996 | | |
| Replicate | 1.563 | | |
| Block x Replicate | 1.367 | | |
| Replicate x Sex | 3.0891 | | |
| Residual | 55.730 | | |

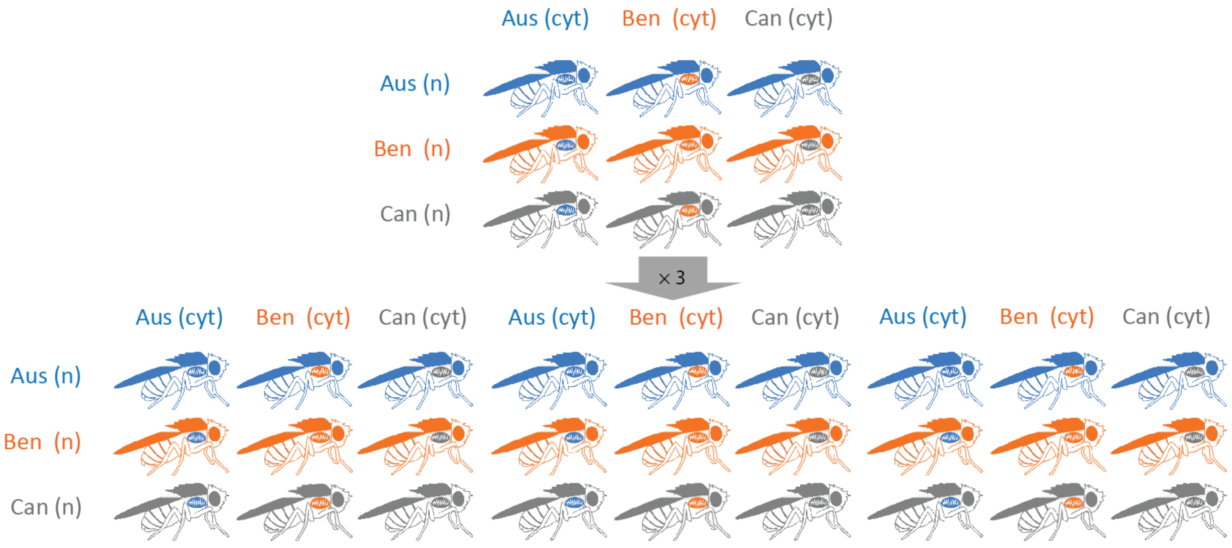

**Supporting Information Figure 1. Creation of the cyto-nuclear lines utilized for this study**. Australian cytoplasm shown in blue, Beninese cytoplasm shown in orange, Canadian cytoplasm shown in grey. Nuclear backgrounds are represented by body coloration, cytoplasms are represented by coloration of the picture of the mitochondrion that sits within each fly. The 3 × 3 cyto-nuclear crossing scheme was triplicated at the outset of their creation to yield 27 individual lines.

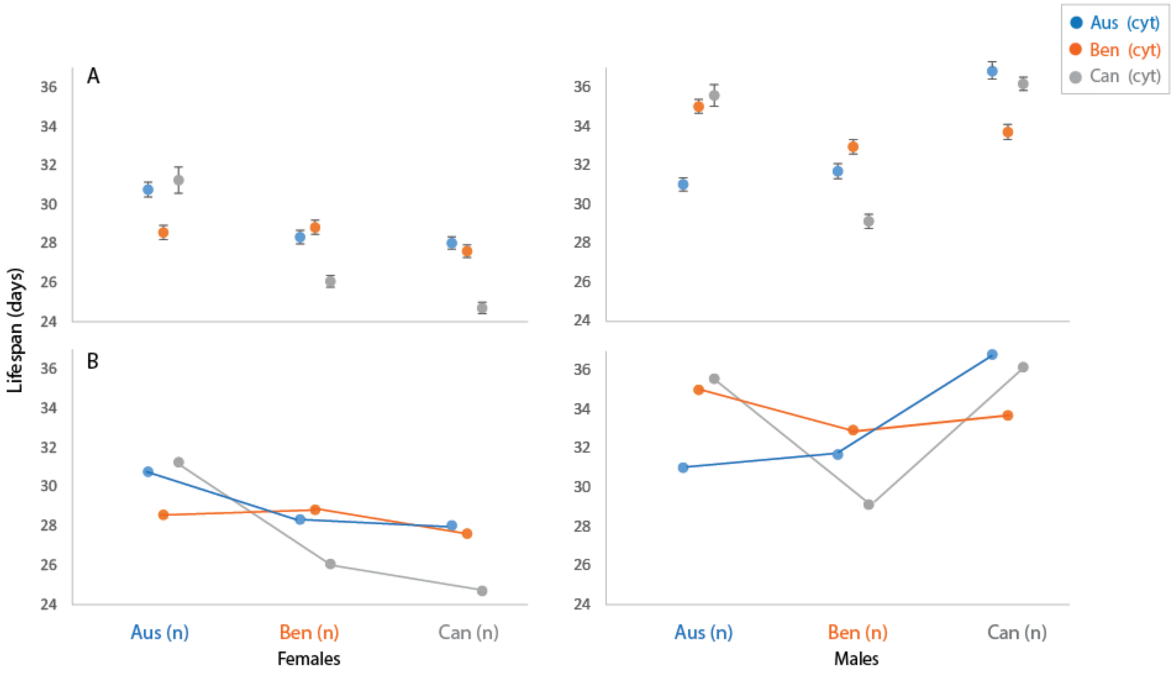

**Supporting Information Figure 2.** Homing in on mitochondrial genetic effects. **A.** Mean longevity of each of the three cytoplasms across the three nuclear backgrounds with the unique mitotypes (AA3, CA1, and CA3) were omitted from the dataset. Each colour denotes a different cytoplasm (blue = Australia, orange = Benin, grey = Canada). Error bars indicate standard error based on calculated means. **B**. Reaction norms of the performance of each of the three cytoplasms across the three nuclear backgrounds (x axis) as measured by mean longevity (y axis). Where the unique mitotypes (AA3, CA1, and CA3) were omitted from the dataset. Australia shown in blue, Benin shown in orange, Canada shown in grey.

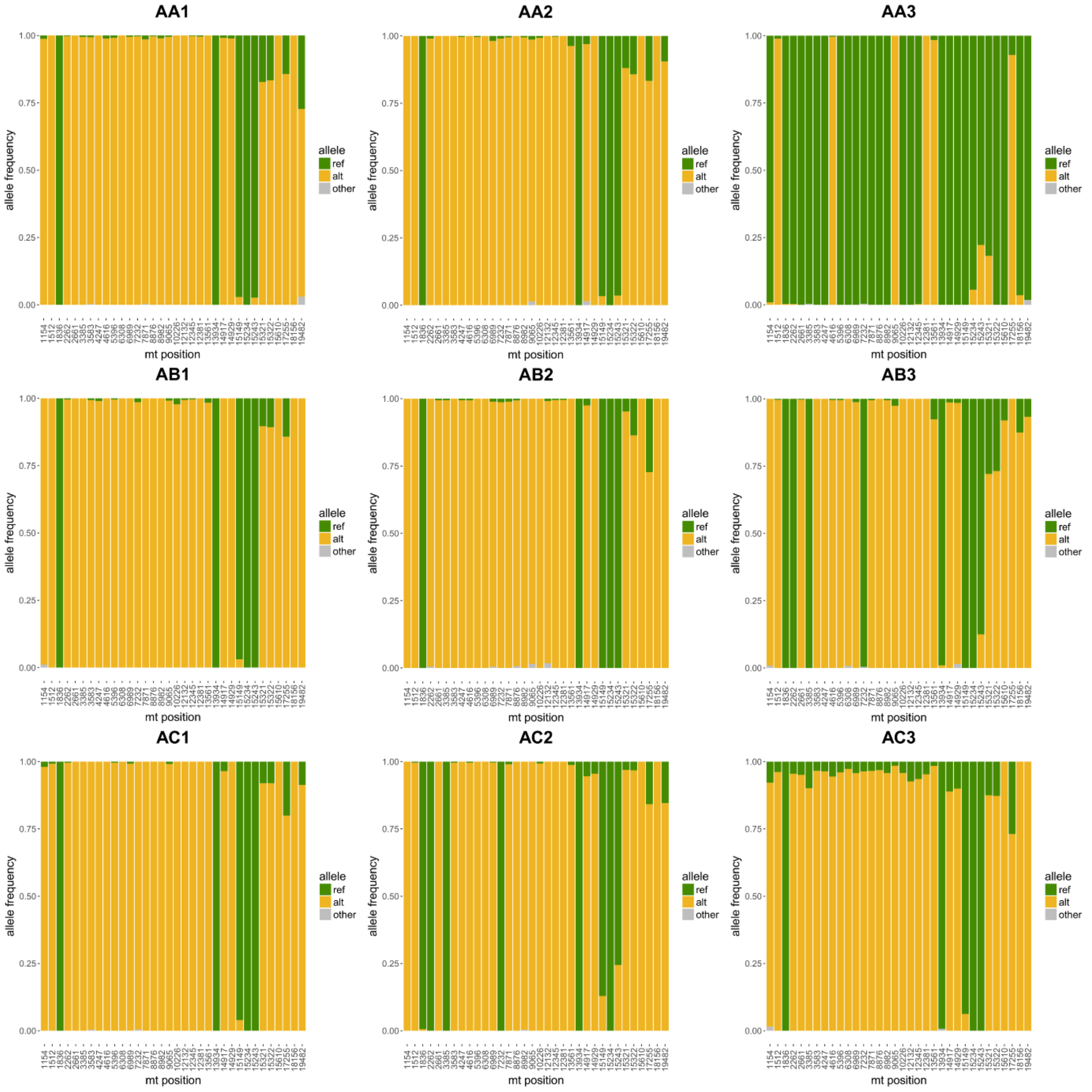

**Supporting Information Figure 3.** Allele frequencies at the 34 significantly differentiated SNP sites in lines with Australian-derived mtDNA. Reference (ref) alleles (green) are according to the mitochondrial reference sequence NC_024511. Alternative (alt) alleles (yellow) correspond to the most frequent non-reference nucleotide at each site. Any other nucleotide states are shown in grey.

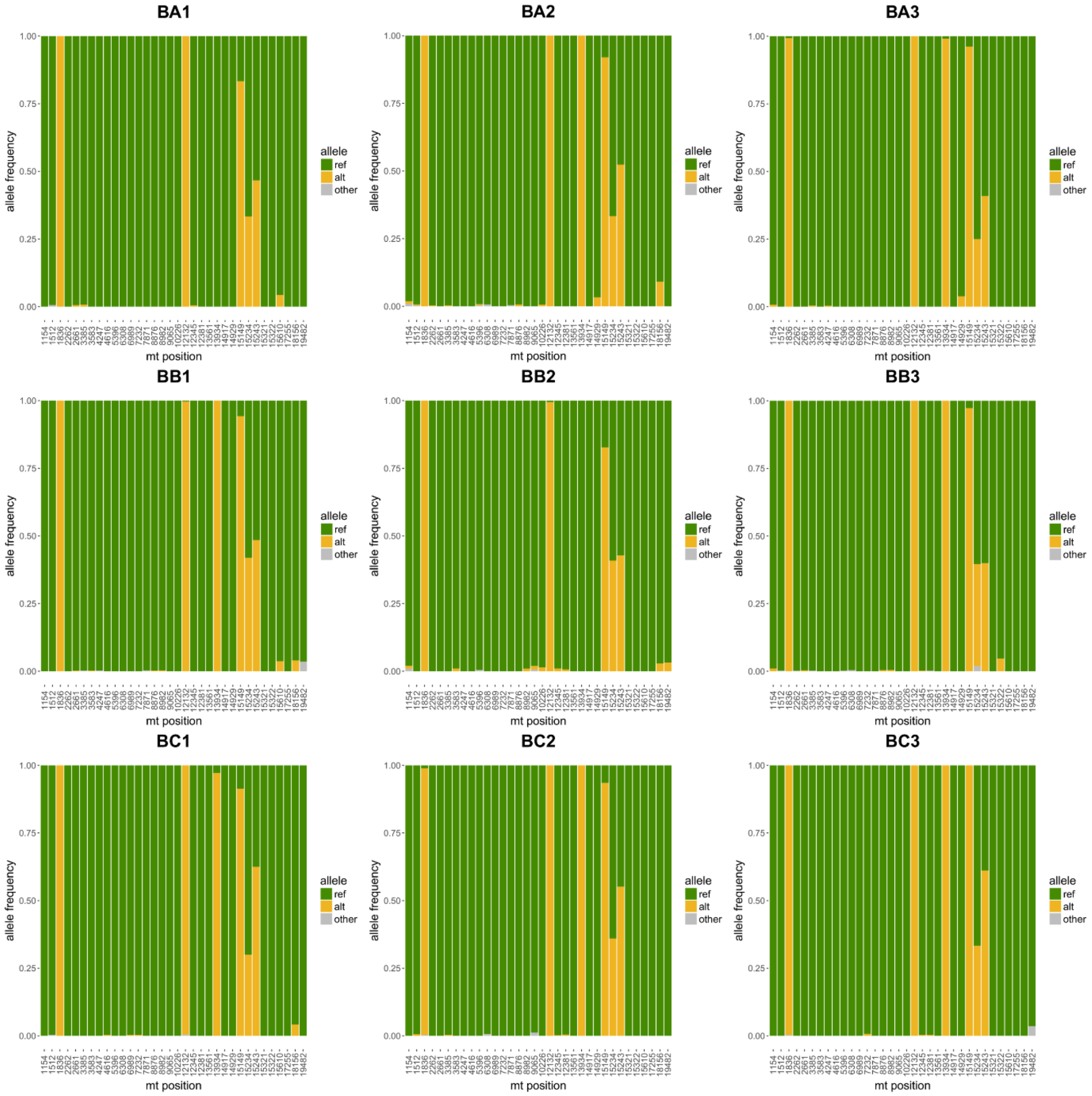

**Supporting Information Figure 4.** Allele frequencies at the 34 significantly differentiated SNP sites in lines with Beninese-derived mtDNA. Reference (ref) alleles (green) are according to the mitochondrial reference sequence NC_024511. Alternative (alt) alleles (yellow) correspond to the most frequent non-reference nucleotide at each site. Any other nucleotide states are shown in grey.

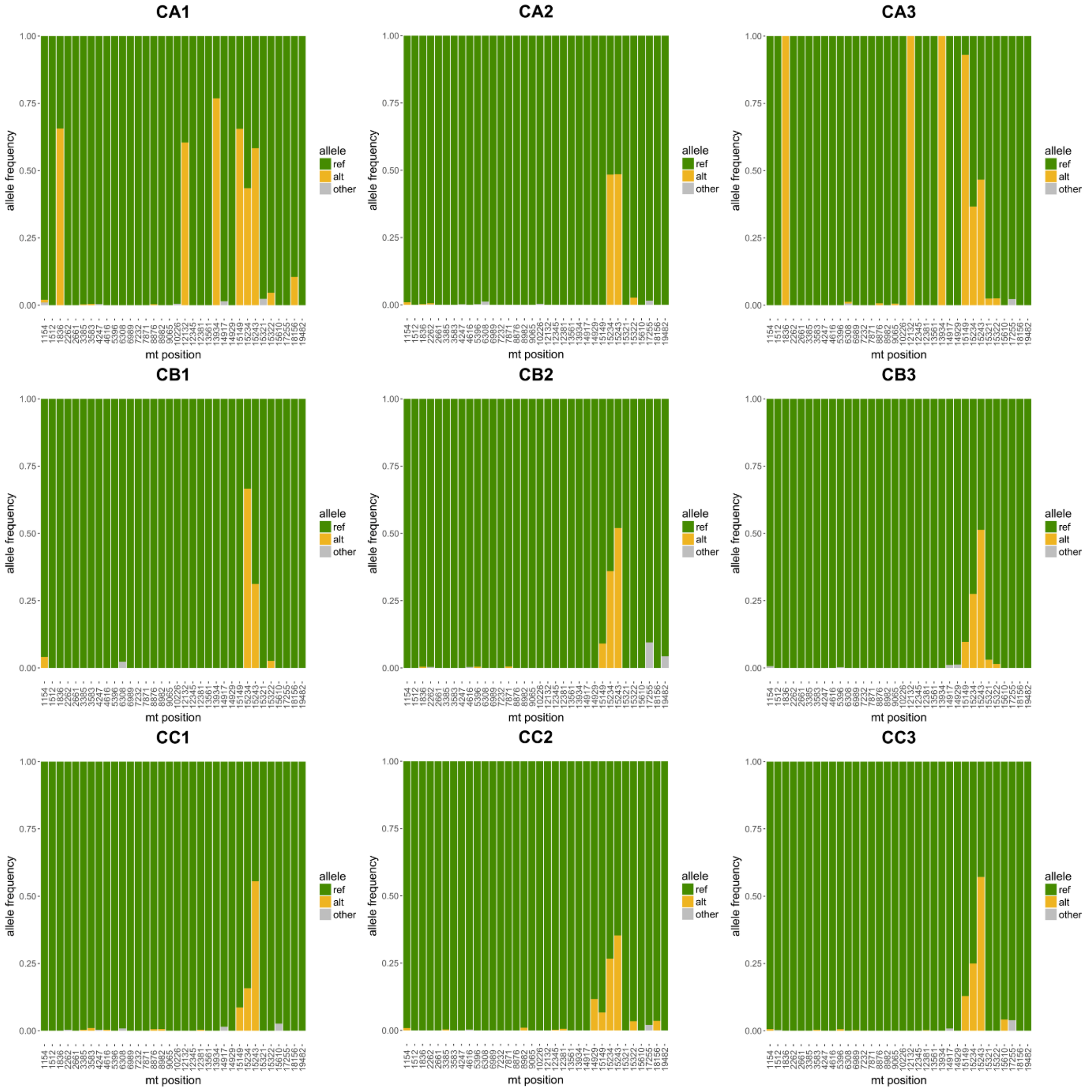

**Supporting Information Figure 5.** Allele frequencies at the 34 significantly differentiated SNP sites in lines with Canadian-derived mtDNA. Reference (ref) alleles (green) are according to the mitochondrial reference sequence NC_024511. Alternative (alt) alleles (yellow) correspond to the most frequent non-reference nucleotide at each site. Any other nucleotide states are shown in grey.

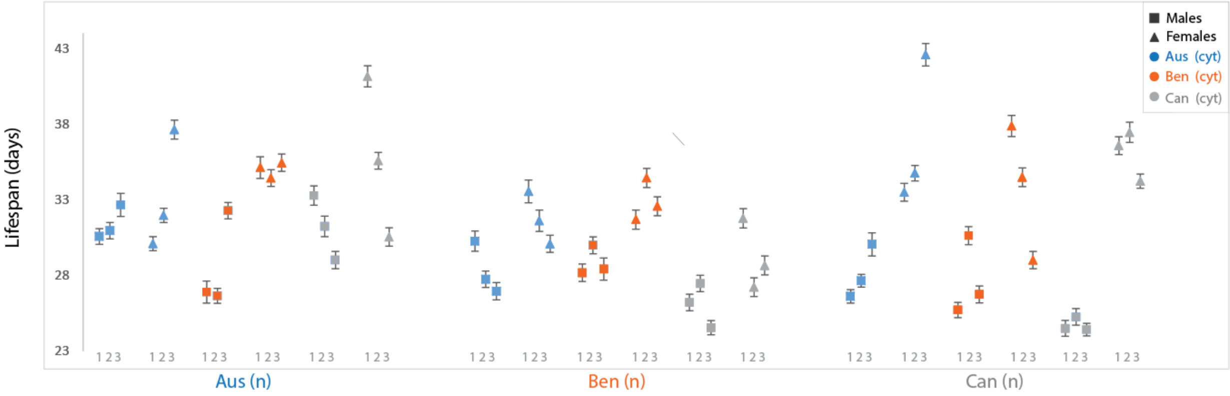

**Supporting Information Figure 6.** Average longevity for all replicates across each of the cyto-nuclear population replicates (54 in total). Data points have been color-coded to reflect the cytoplasm each lines carries. The Australian cytoplasms are shown in blue. Beninese cytoplasms are shown in orange, Canadian cytoplasms shown in grey. Biological replicates are represented in order (triplicate 1, 2, and 3) with respect to the x-axis as indicated by the grey numbers. Error bars indicate standard error.

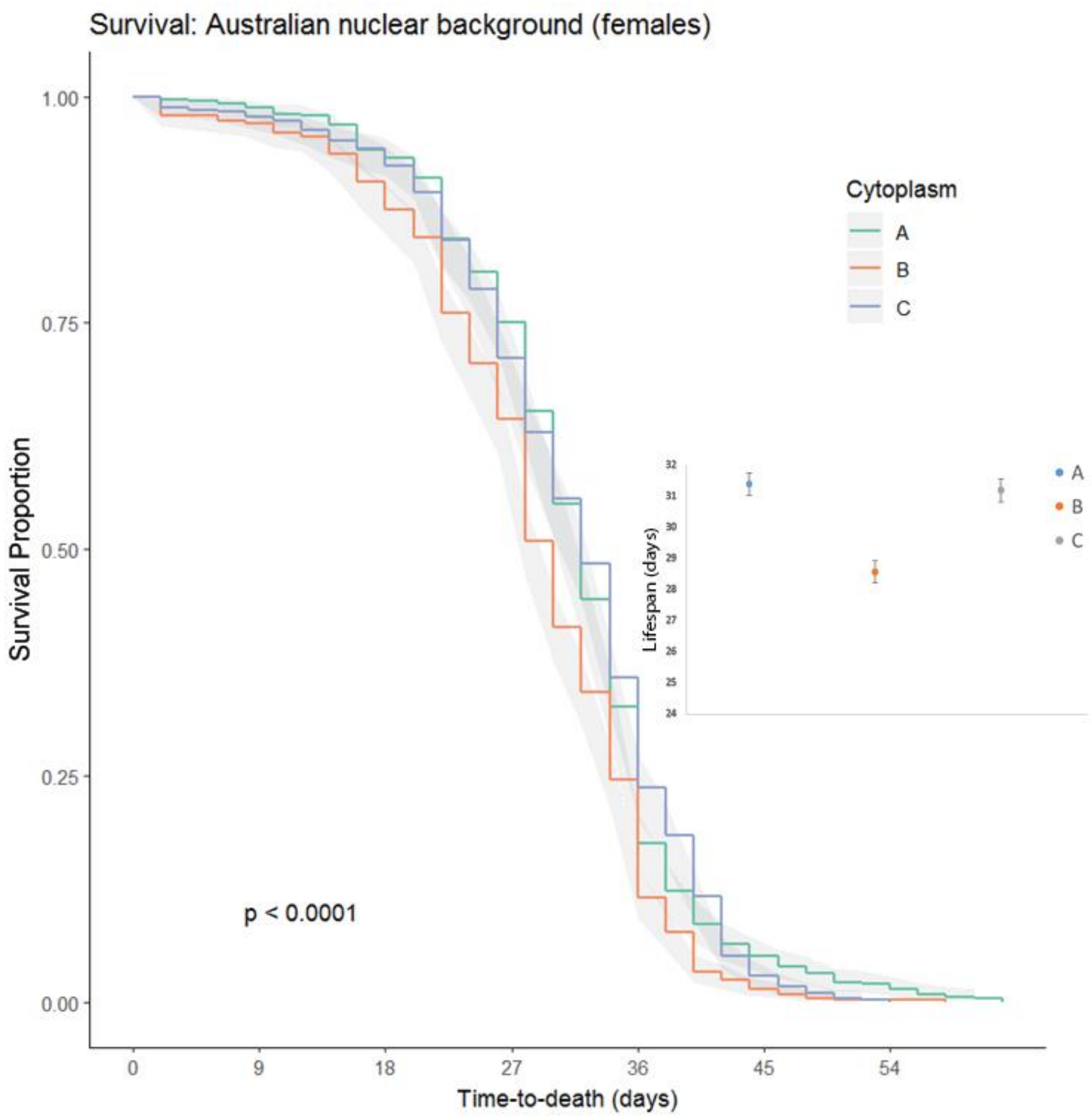

**Supporting Information Figure 7.** Partitioning of cytoplasmic and nuclear sources of variance. Cytoplasmic effects of genotypes survival (± 95% C.I.), for females with the Australian nuclear background only. Each colour denotes a different cytoplasm: Australia shown in green, Benin shown in orange, Canada shown in purple. Mean lifespan (± 1 standard error) of each of the three cytoplasms expressed against each of the same nuclear background is shown in the right-hand panel. Lifespan (as measured in days) is indicated on the vertical axis, with each cytoplasm on the horizontal axis. Each colour denotes a different cytoplasm: Australia shown in blue, Benin shown in orange, Canada shown in grey.

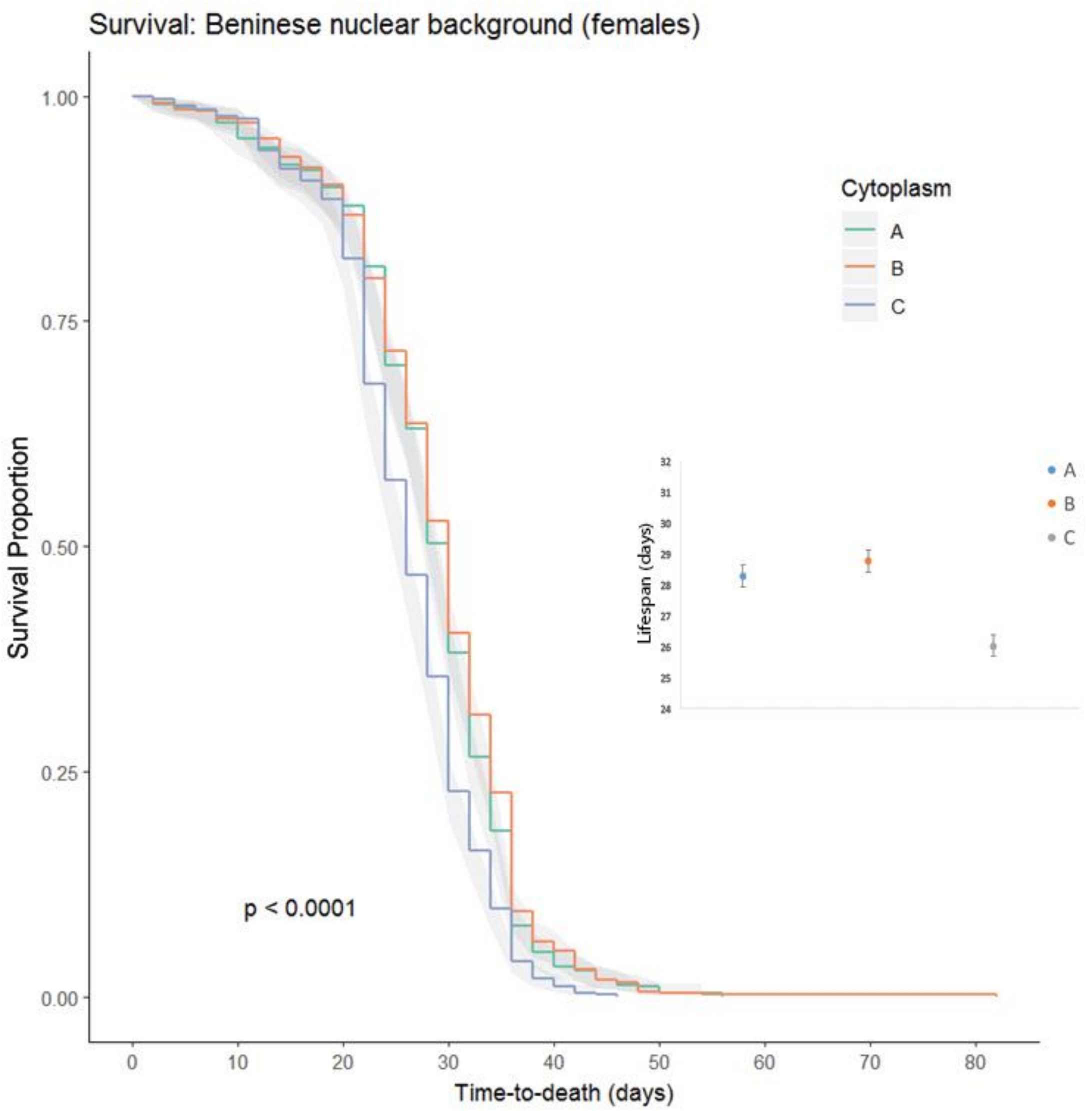

**Supporting Information Figure 8.** Partitioning of cytoplasmic and nuclear sources of variance. Cytoplasmic effects of genotypes survival (± 95% C.I.), for females with the Beninese nuclear background only. Each colour denotes a different cytoplasm: Australia shown in green, Benin shown in orange, Canada shown in purple. Mean lifespan (± 1 standard error) of each of the three cytoplasms expressed against each of the same nuclear background is shown in the right-hand panel. Lifespan (as measured in days) is indicated on the vertical axis, with each cytoplasm on the horizontal axis. Each colour denotes a different cytoplasm: Australia shown in blue, Benin shown in orange, Canada shown in grey.

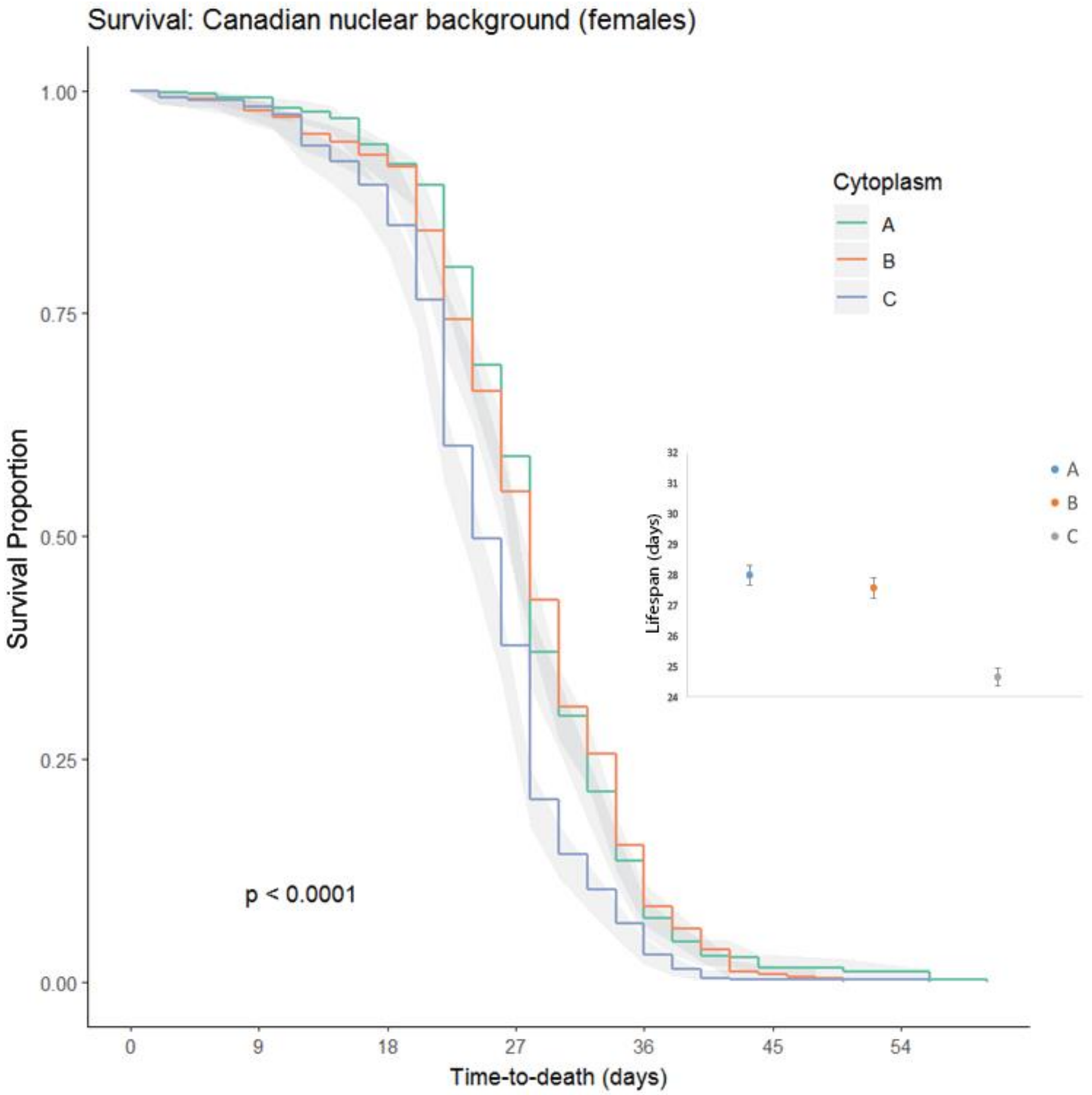

**Supporting Information Figure 9.** Partitioning of cytoplasmic and nuclear sources of variance. Cytoplasmic effects of genotypes survival (± 95% C.I.), for females with the Canadian nuclear background only. Each colour denotes a different cytoplasm: Australia shown in green, Benin shown in orange, Canada shown in purple. Mean lifespan (± 1 standard error) of each of the three cytoplasms expressed against each of the same nuclear background is shown in the right-hand panel. Lifespan (as measured in days) is indicated on the vertical axis, with each cytoplasm on the horizontal axis. Each colour denotes a different cytoplasm: Australia shown in blue, Benin shown in orange, Canada shown in grey.

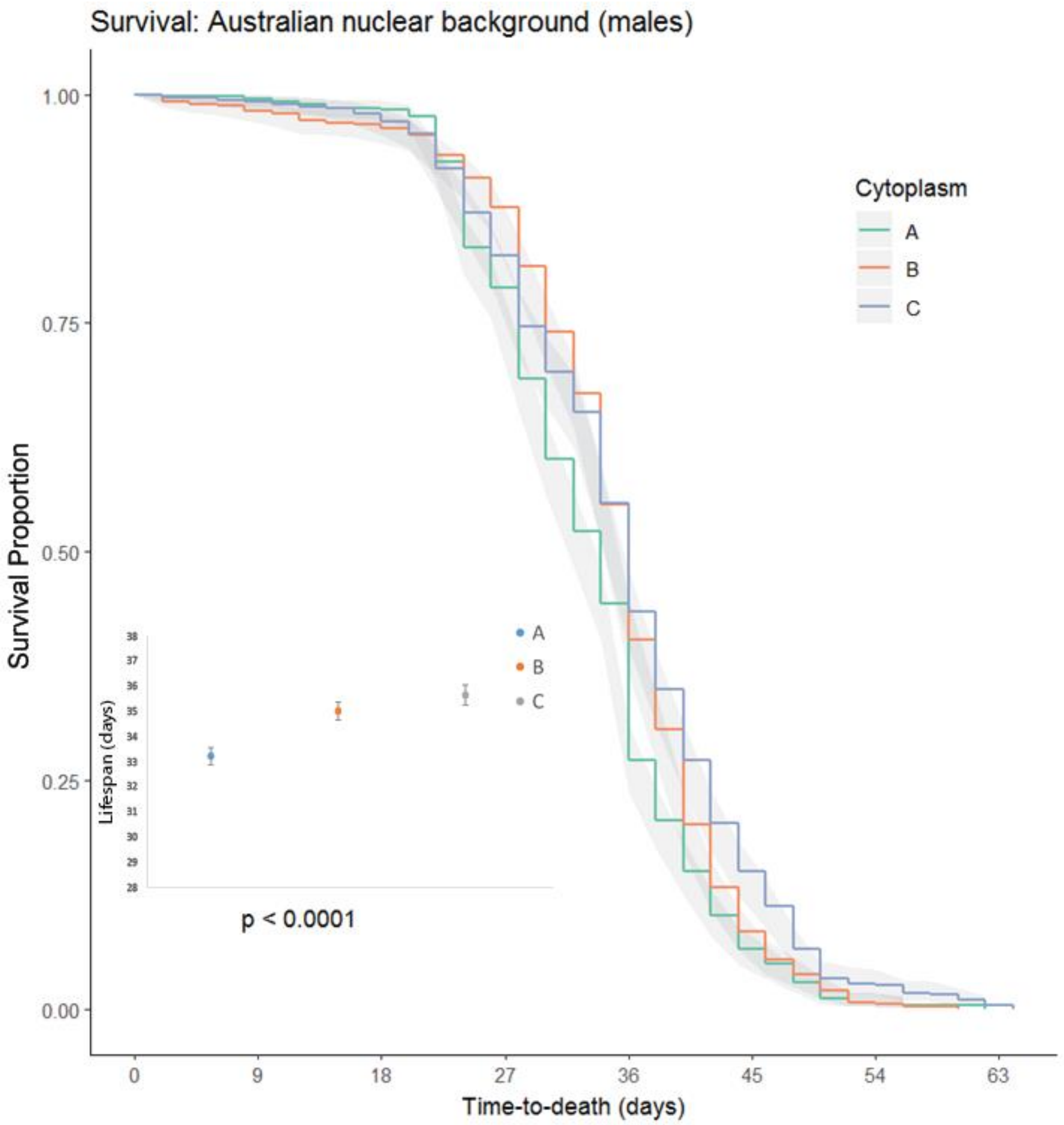

**Supporting Information Figure 10.** Partitioning of cytoplasmic and nuclear sources of variance. Cytoplasmic effects of genotypes survival (± 95% C.I.), for males with the Australian nuclear background only. Each colour denotes a different cytoplasm: Australia shown in green, Benin shown in orange, Canada shown in purple. Mean lifespan (± 1 standard error) of each of the three cytoplasms expressed against each of the same nuclear background is shown in the left-hand panel. Lifespan (as measured in days) is indicated on the vertical axis, with each cytoplasm on the horizontal axis. Each colour denotes a different cytoplasm: Australia shown in blue, Benin shown in orange, Canada shown in grey.

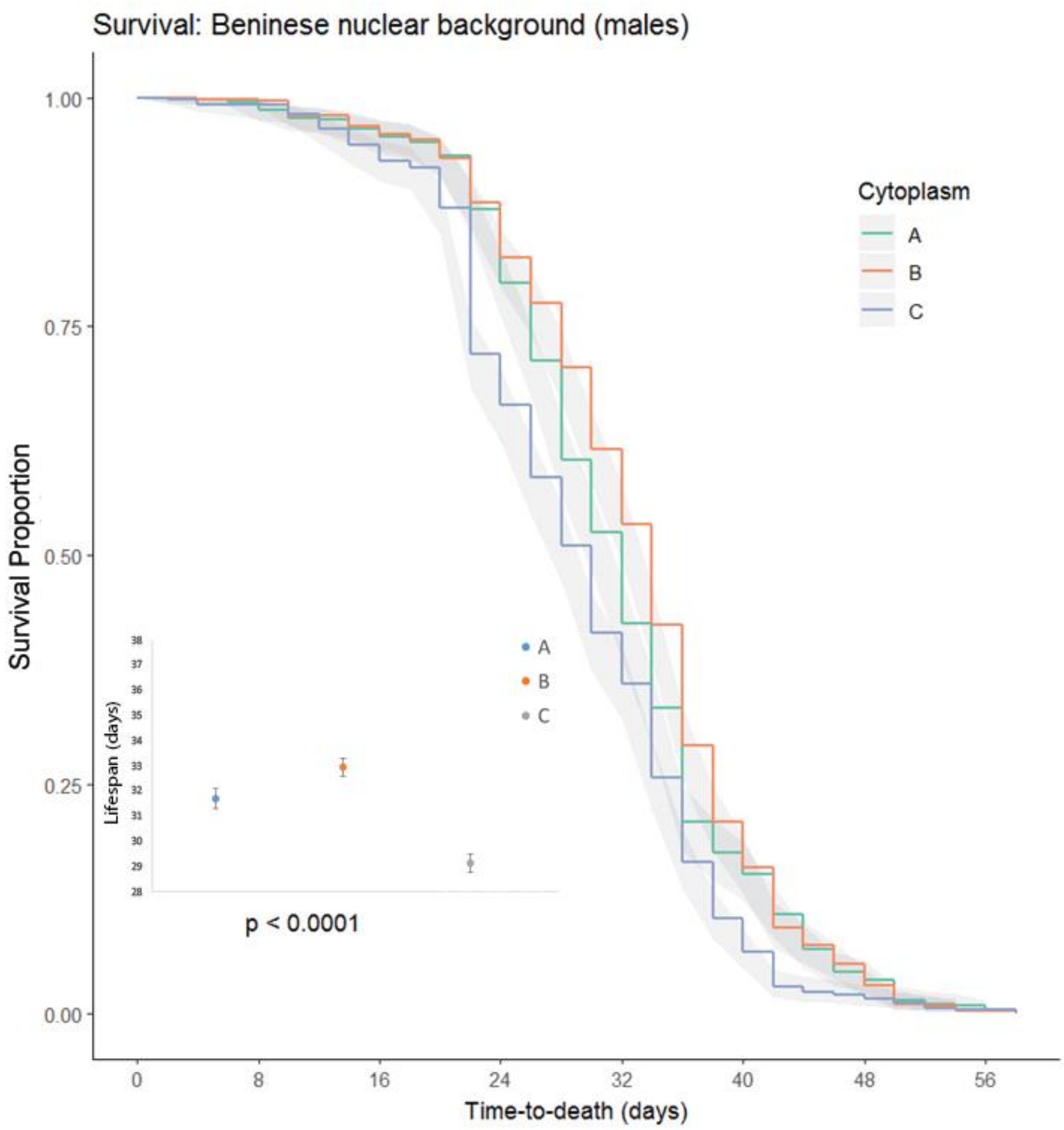

**Supporting Information Figure 11.** Partitioning of cytoplasmic and nuclear sources of variance. Cytoplasmic effects of genotypes survival (± 95% C.I.), for males with the Beninese nuclear background only. Each colour denotes a different cytoplasm: Australia shown in green, Benin shown in orange, Canada shown in purple. Mean lifespan (± 1 standard error) of each of the three cytoplasms expressed against each of the same nuclear background is shown in the left-hand panel. Lifespan (as measured in days) is indicated on the vertical axis, with each cytoplasm on the horizontal axis. Each colour denotes a different cytoplasm: Australia shown in blue, Benin shown in orange, Canada shown in grey.

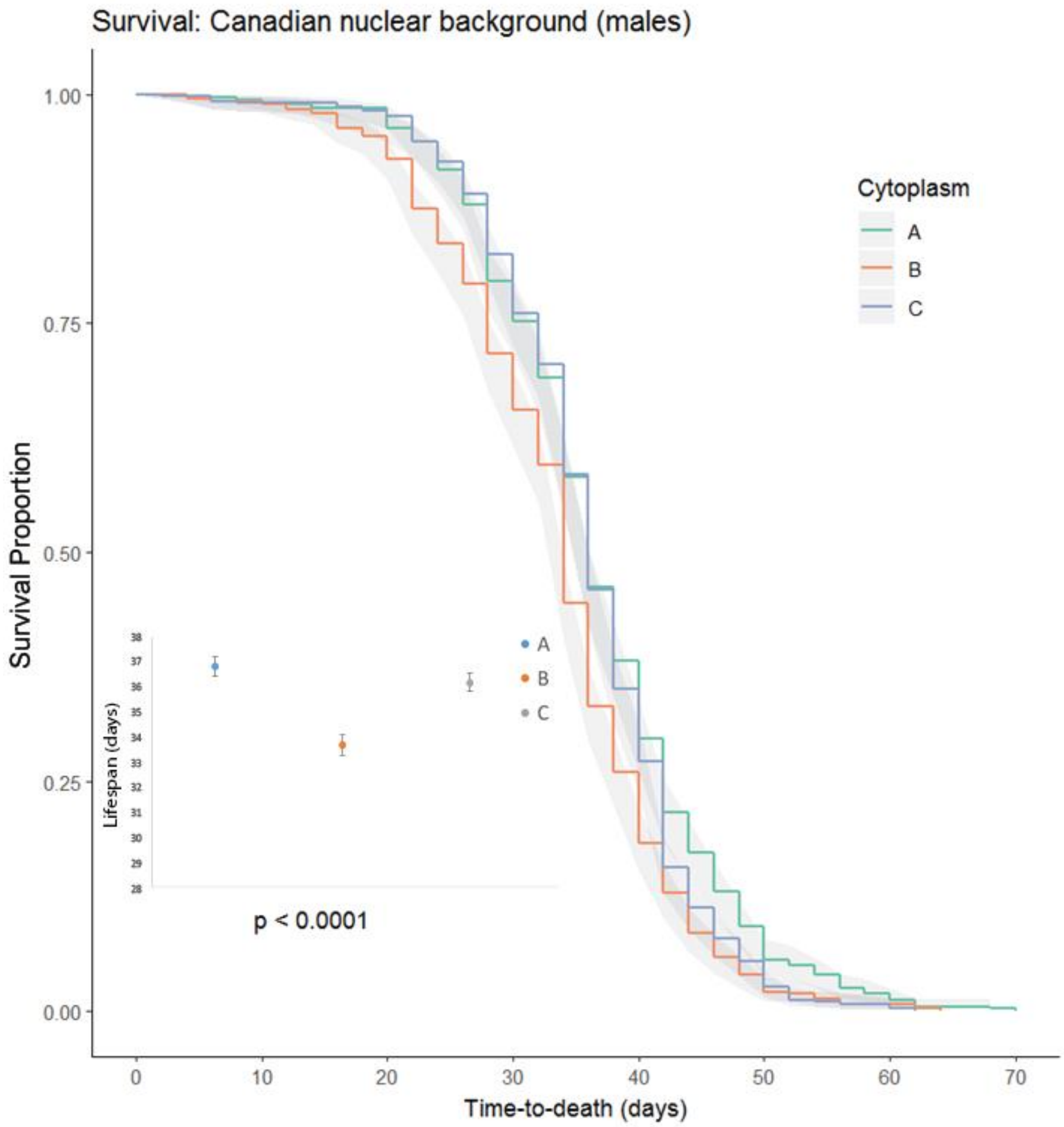

**Supporting Information Figure 12.** Partitioning of cytoplasmic and nuclear sources of variance. Cytoplasmic effects of genotypes survival (± 95% C.I.), for males with the Canadian nuclear background only. Each colour denotes a different cytoplasm: Australia shown in green, Benin shown in orange, Canada shown in purple. Mean lifespan (± 1 standard error) of each of the three cytoplasms expressed against each of the same nuclear background is shown in the left-hand panel. Lifespan (as measured in days) is indicated on the vertical axis, with each cytoplasm on the horizontal axis. Each colour denotes a different cytoplasm: Australia shown in blue, Benin shown in orange, Canada shown in grey.
